# Evolution and Diversification of the Aposematic Poison Frog, *Oophaga pumilio,* in Bocas del Toro

**DOI:** 10.1101/2024.08.02.606438

**Authors:** Diana Aguilar-Gómez, Layla Freeborn, Lin Yuan, Lydia L. Smith, Alex Guzman, Andrew H. Vaughn, Emma Steigerwald, Adam Stuckert, Yusan Yang, Tyler Linderoth, Matthew MacManes, Corinne Richards-Zawacki, Rasmus Nielsen

## Abstract

The aposematic strawberry poison frog, *Oophaga pumilio*, is an iconic model system for studying the evolution and maintenance of color variation. Through most of its range, this frog is red with blue limbs. However, frogs from the Bocas del Toro Province, Panama, show striking variance in color and pattern, both sympatrically and allopatrically. This observation contradicts standard models of the evolution of aposematism and has led to substantial speculation about its evolutionary and molecular causes. Since the enigma of *O. pumilio* phenotypic variation is partly unresolved because of its large, ∼ 6.7 Gb genome, we here sequence exomes from 347 individuals from ten populations and map a number of genetic factors responsible for the color and pattern variation. The *kit* gene is the primary candidate underlying the blue-red polymorphism in Dolphin Bay, where an increase in melanosomes is correlated with blue coloration. Additionally, the *ttc39b* gene, a known enhancer of yellow-to-red carotenoid conversion in birds, is the primary factor behind the yellow-red polymorphism in the Bastimentos West area. The causal genetic regions show evidence of selective sweeps acting locally to spread the rare phenotype. Our analyses suggest an evolutionary model in which selection is driving the formation of new morphs in a dynamic system resulting from a trade-off between predation avoidance, intraspecific competition, and mate choice.

## Introduction

The strawberry poison frog, *Oophaga pumilio*, has bright red coloration in most of its continental distribution ^1^. This coloration is essential in aposematism, a warning signal to predators ^2^. However, in the Bocas del Toro Province of Panama, this species exhibits remarkable variation in color and pattern ^3^. Colors on the islands include yellow, orange, red, green, blue, and some intermediate phenotypes (Fig. 1A). On top of the color variation, there is also pattern diversity, with some frogs being a uniform color, others having speckles, and some having large spots or bands of black. The formation of the archipelago was recent–1000 to 9000 years ago ^4^–suggesting that the dramatic phenotypic differences emerged in a very short period of time. Most color variation is allopatric between islands (polytypism) ^5^, with each separate island population being monomorphic. However, a few islands have sympatric variation (polymorphism) ^6^. This variation has made *O. pumilio* a model for understanding the mechanisms underlying the maintenance of phenotypic variation, with hundreds of studies on *O. pumilio* focused on ecology ^3,7–11^, toxicity ^1,12^, sexual selection ^13–15^, natural selection ^10,16^ and population history ^17^.

**Fig. 1.**
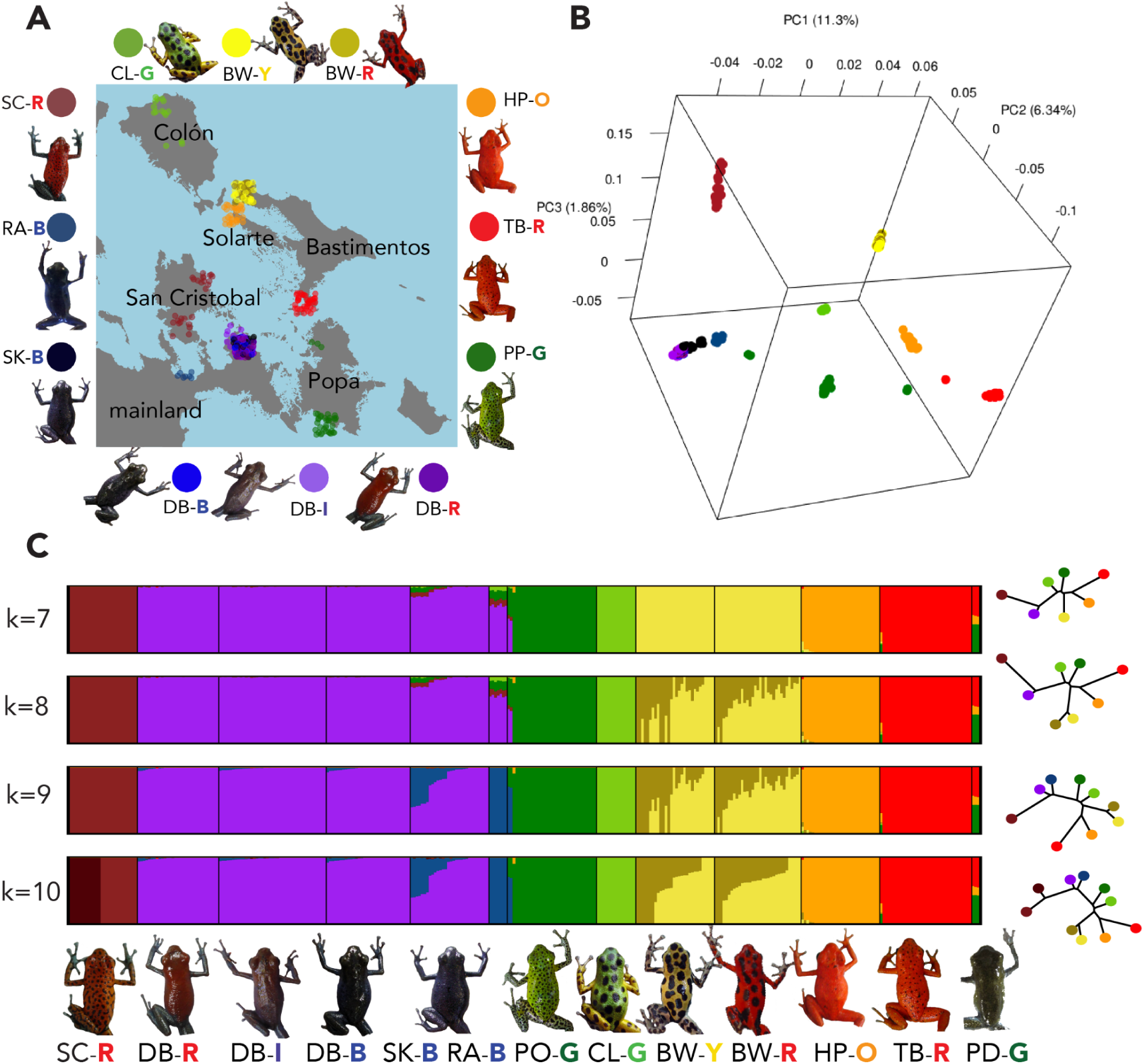
Population structure of *Oophaga pumilio* in Bocas del Toro Archipelago. A) Sampling locations B) PCA of genome-wide variation C) Structure plots and trees using k=7 to k=10 ancestry components. Labels of populations have an added letter to specify the color of the individuals (R-red, I-intermediate, B-blue, G-green, Y-yellow, and O-orange). Popa Island (PP) individuals split into South Popa (PO-G) and North Popa (PD-G)

Standard theory on aposematism suggests that natural selection acts to reduce variation ^2,18^, making the high degree of phenotypic variability observed in Bocas del Toro somewhat enigmatic. There are multiple hypotheses regarding the emergence and maintenance of this variation. Some of them include a) viability selection, where color variation may have evolved as an adaptation to different environments or predators as an honest signal of toxicity ^7,12,19^; b) sexual selection, in the form of female mate choice and male-male competition ^6,14,2017,216,14,20^; c) geographic isolation, since most morphs are restricted to different islands ^17,21^; d) and a lack of predators combined with neutral drift ^22^. Many of the hypotheses can be combined, with most authors agreeing on a combination of several factors likely responsible for this variation ^3,17,21^. *Oophaga pumilio* has complex parental care and tadpoles learn to recognize traits of their parents (i.e., imprinting), which may influence their own mating behavior later in life ^6^. *Oophaga pumilio* shows evidence of assortative mate preferences whereby females ^23^ of allopatric populations prefer to mate with males of their own morph ^24,25^. Additionally, males compete by defending territories and are generally more aggressive towards males of their own morph, leading to the possibility of sexual selection favoring rare morphs ^13,20^. Imprinting, female bias and male bias together are the main reasons why sexual selection is often suggested to drive the maintenance and divergence of morphs ^6^.

Moreover, the genetic and physiological mechanisms underlying such phenotypic variation are becoming increasingly understood. Advancements in the cell biology of color variation in vertebrates have shed light on the role of specialized pigment cells, known as chromatophores, in producing and manipulating colors and patterns on the skin ^26–29^. There are several types of chromatophores, including melanophores, which contain black/brown pigments; iridophores, which contain reflective platelets; and xanthophores, which contain yellow/orange; and erythrophores, which contain red pigments ^26,27^. These cells are arranged in layers in the skin, and their contraction or expansion can change the size and distribution of the pigments, resulting in changes in color and pattern ^30^. For example, xanthophores have a filtering function and are associated with yellow, orange and red phenotypes. Iridophores scatter light, reflecting shorter wavelengths of light, associated with blue/green phenotypes and melanophores absorb longer wavelengths that are not scattered by iridophores ^28^.

Many of these described genetic and physiological mechanisms create the vast array of colors and patterns across amphibian diversity. The ancestral coloration in *O. pumilio* is red ^31^, but there are also green and blue morphs in Bocas del Toro. Amphibians do not have chromatophores that produce green or blue pigments. Instead, blue can be produced as a structural color due to diffraction or interference of light ^32^, caused by specific layerings of different chromatophores ^28,30^. In other frogs, green coloration is then produced by combining yellow pigments with blue structural coloration. For example, in the American green tree frog, *Hyla cinerea*; the combined effect of xanthophores and iridophores forms green ^30^; iridophores filter the light selectively to create blue coloration, then an overlaying layer of yellow xanthophores results in green ^28,30^. Carotenoids primarily located in the xanthophores are involved in yellow/orange/red coloration but are also crucial for green coloration in frogs. For example, a higher carotenoid intake diet results in much brighter green coloration in some frogs ^33,34^. Alternatively, the bones or lymph of some frogs are tinted through a mechanism called chlorosis, defined as an unusually high concentration of the pigment biliverdin ^35^. However, dendrobatid frogs such as *O. pumilio* do not appear to have elevated biliverdin levels ^35,36^ and thus the blue/green coloration in some morphs is thought to be at least somewhat structural.

Previous genetic research on *O. pumilio* has been restricted to mitochondrial DNA, microsatellite ^37^, amplified fragment length polymorphism (AFLP) ^10^, differential expression ^38,39^ studies, and RADSeq ^40^, in addition to the generation of a reference genome ^41^. However, population-scale genomic data for this species has been slow in coming due to the large (∼6.76 Gb) and highly repetitive genome of *O. pumilio,* which makes this type of approach technically challenging and expensive. To address these challenges, we present exome sequencing data from 347 individuals from Bocas del Toro. We use GWAS coupled with selection scans to identify genes underlying the color polymorphisms and use further population genetic analyses of these genes to answer decades-old questions regarding the causes of the Bocas del Toro color variation.

## Results

### Population Structure and Demographic History in Bocas del Toro

To infer the population structure and history of the Bocas del Toro *O. pumilio*, we sampled 347 *Oophaga pumilio* frogs from eight different morphs, each defined by a unique combination of patterning and coloration: Aguacate (blue, two sampling locations: Rana Azul (RA-B) and Shark Hole (SK-B)), Dolphin Bay (DB, B blue, I intermediate, and R red), Bastimentos West (BW, R red and Y yellow), Tranquilo Bay (TB-R, red), Hospital Point (HP-O, orange), Popa (PP-G, green), Colón (CL-G, green) and San Cristobal (SC-R, red) (Fig. 1A). We performed Principal Component Analysis (PCA; Fig. 1B) on 10,788,793 filtered SNPs. PC1 separates frogs geographically from east to west, except PP-G, which is placed between TB-R and mainland morphs from the peninsula (SK, DB and RA). PC2 correlates with a north-south axis of Bastimentos Island (BW and TB). PC3 separates the SC-R morph from the mainland frogs from the peninsula.

We used OHANA^42^ for unsupervised ancestry component assignment, testing for k=7 to k=10. Most sampled populations are allocated to their own component using k=7 (Fig. 1C), except for the mainland DB and AG, which share a component. The Aguacate morph separates into Rana Azul (RA-B) and Shark Hole (SK-B) subpopulations at >k=9. Some SK frogs are admixed between the DB and RA. Admixture analyses did not separate polymorphic populations BW (yellow-red) and DB (blue-red) into different ancestry components, even up to k=10 components (Fig. 1B and C). Three PP-G morph individuals sampled in the north of Popa Island are estimated to be admixed with the TB population and separate from other PP-G individuals in PCA (Fig. 1B). Therefore, we split the Popa morph into South Popa (PO-G) and North Popa (PD-G) for subsequent analyses.

We calculated genome-wide pairwise F_ST_ (Sup Table 1). Allopatric populations showed trends similar to previous studies using microsatellites and amplified fragment length polymorphisms ^10,17^. Sympatric morphs within the same population (BW and DB) that could not be separated in the structure analysis or PCA, have low values of F_ST_ (∼0.01), and are indistinguishable using an average genomic tree (Sup Fig. 1). Some populations within the same island are genetically differentiated. For example, TB and BW from Bastimentos Island show no admixture and are more diverged genome-wide (F_ST_ >0.2) than populations from different islands. The North Popa population (PD) is less diverged from the red TB population from Bastimentos (F_ST_ =0.126) than from the southern Popa population, PO (F_ST_ =0.133). Despite their genetic differences, both Popa Island populations are phenotypically indistinguishable.

Previous studies detected no evidence of migration in Bocas del Toro ^17^. However, using AdmixtureBayes^43^ we tested topologies with up to four admixture events (Sup Fig. 2). Topologies allowing less than four migration events can be rejected by 4-population (ABBA/BABA) tests (Sup Table 2), providing evidence for substantial migration among islands. For example, the topology recovered with 100% probability when specifying zero admixture events, has the subgraph ((TB, PD)PO), *O. sylvatica*), while the 4-population test shows ((PO,PD)TB), *O. sylvatica*) as the most prevalent topology. The top three topologies identified by AdmixtureBayes^43^ represent 74% of the probability with four admixture events. Interpreting these topologies regarding colonization scenarios, they suggest an ancestor from the mainland splitting into two lineages, one entering the archipelago from the north through Colon Island. Another from the south via the Aguacate Peninsula (Sup Fig. 2). The northern lineage subsequently splits into (HP, BW) and (TB, PD). The southern lineage is (RA, SC, (DB, SK)) with admixture from RA to SK (seen in structure analysis, Fig. 1C). CL is admixed between north and south lineages, its proximity to the mainland likely facilitating migration. Similarly, PO is admixed between PD and mainland ancestry. Popa Island split from Aguacate Peninsula ∼1,000 years ago^4^ and is currently separated by a small stretch of water (Sup Figure 2). All three admixture topologies indicate bidirectional migration between Aguacate Peninsula and Popa Island. Our results suggest extensive gene flow between Bocas del Toro populations, contradicting a simple model of population splits without subsequent gene flow. Multiple colonization waves during past glaciation eras^4^ and/or continued ongoing gene-flow likely explain the observed patterns.

To further investigate the time-scale of migration and the split between populations, we performed demographic inference for pairs of populations using dadi ^44^. Assuming a mutation rate of 10^-9 45,46^ the divergence time between populations is close to the island formation times (Sup Table 3). For example, Colon Island and Bastimentos Island separated ∼5,200 years ago ^4^. Our results suggest the split time between the two closest (genetically and geographically) populations CL-BW was ∼8,000 years ago (Sup Table 3).

### Genomic basis of color polymorphism

#### Blue

To uncover the genomic basis of blue coloration, we sampled blue frogs from Rana Azul (RA-B), Dolphin Bay (DB-B), and Shark Hole (SK-B). Since Dolphin Bay is a polymorphic blue population, also harboring red and intermediate phenotypes, we performed a population branch statistic (PBS) scan of DB-B and DB-R frogs, using the TB-R population as an outgroup. The most differentiated gene in DB-B frogs is *kit* (Fig. 2A, B). We also performed a genomic association with spectrometric measurements of blue wavelength (Fig. 2D, E), correcting for population structure using a relatedness matrix. Again, the gene with the strongest association was *kit*. We identified two non-synonymous mutations in *kit*: G959R and V533I. G959R, located in the C-terminal domain of Kit (Fig. 2J), is fixed for the ancestral allele fixed in bright populations such as TB (red), BW (yellow/red), and HP (orange), but instead segregates in DB and in duller/darker colored populations (Fig. 2F). Meanwhile, V533I is located in the transmembrane domain of kit. This mutation is fixed for the ancestral allele in BW and has a higher frequency in darker populations (Fig. 2F). Promisingly, a previous expression study also identified mutations unique to blue frogs in the kit gene ^38^.

**Fig. 2.**
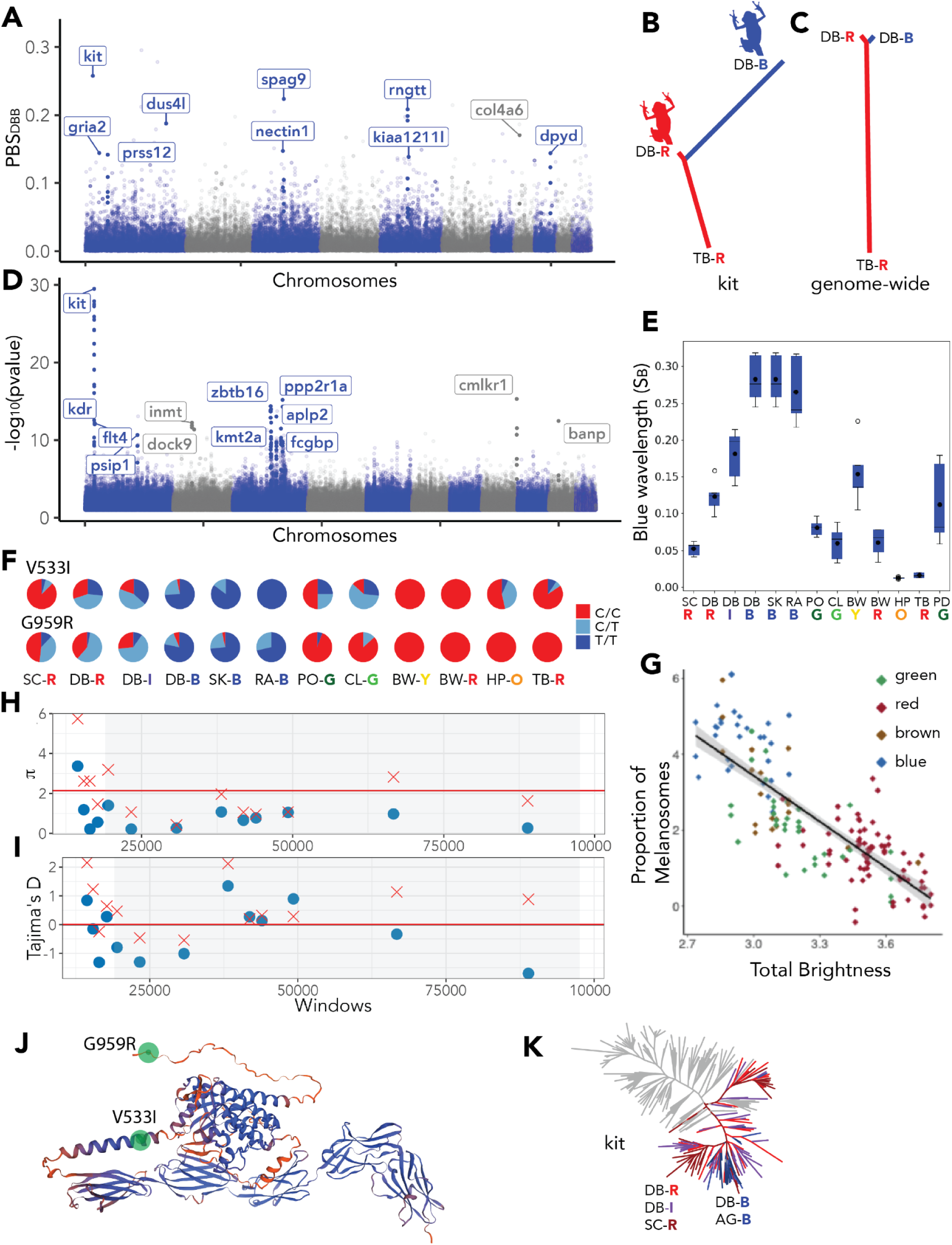
Genetic basis of blue coloration in *O.pumilio.* A) PBS scan of Dolphin Bay Blue (DB-B) branch against DB-R and TB-R. Mapped to *O.pumilio*, plot using synteny, alternating colors are chromosomes from *O.sylvatica* B) PBS tree of *kit* scaffold C) PBS genome-wide D) Genomic association to blue color wavelengths. E) Blue wavelength (SB) measurements grouped by population F) Allele frequency of non-synonymous mutations of *kit.* G) Proportion of melanosomes in different color morphs of *O. pumilio*, reproduced from Freeborn (2020). H) Pairwise differences in 1000 sites windows in the *kit* scaffold. The shaded area contains the protein-coding region. Blue dots correspond to DB-B and red crosses to DB-R. The red and black lines represent the genome-wide average in DB-R and DB-B, respectively I) Same format as the previous plot with Tajima’s D. J) Structure of kit in *O.pumilio*, non-synonymous mutations highlighted in green. K) gene tree of *kit,* mainland populations, and San Cristobal are colored according to the dorsal coloration of the frog, all the other island individuals in gray.

To investigate selection acting on *kit*, we compared local values of average pairwise differences (π) and Tajima’s D to genome-wide values in DB. The blue morph has reduced variability and Tajima’s D, which suggests positive selection in this morph (Fig. 2H, I). We also performed a HKA test to determine if the ratio of polymorphism to divergence in our candidate genes was consistent with genome-wide values. DB-B frogs have relatively fewer polymorphic sites in *kit* than DB-R frogs. SK, the monomorphic blue population, has significantly fewer polymorphic sites in *kit* (qvalue < 0.05), suggesting a selective sweep (Sup. Table 4).

*Kit* is a tyrosinase kinase receptor involved in melanocyte precursor expansion, survival, and migration ^47^. Whereas iridophores spatially close to melanophores are blue-green, isolated iridophores transmit longer wavelengths such as red ^48^. Interestingly, we found that blue *O. pumilio* have more melanosomes than red and green frogs (Fig. 2G). An alteration in melanosome composition in the frogs’ skin driven by *kit* could explain the change of chromatophore content, resulting in structural blue. The *kit* allele found in blue frogs likely promotes more melanin production, increasing the number of melanophores in blue frogs and making them appear darker. The brighter populations, such as BW, where *kit* is fixed for the reference allele, have fewer melanosomes (Fig. 2G).

In the genome-wide tree, individuals from the DB polymorphic population do not cluster by color (Sup Fig. 1). However, in the *kit* gene tree, all of the blue frogs from DB cluster with the remaining blue frogs from the Aguacate Peninsula, in a clade distinct from the red frogs from DB (Fig. 2K).

#### Yellow, red, and orange

Vertebrates cannot produce carotenoids, which are used to make yellow, red, and orange pigments, and instead obtain them through their diet ^29^. The only yellow morph in Bocas del Toro is in Bastimentos West (BW-Y). There is very little differentiation genome-wide between yellow and red color morphs on Bastimentos (genome-wide F_ST_ of ∼0.01, Sup Table 1). However, a few genes have very different allele frequencies between morphs. Some SNPs have an F_ST_ value of 0.98 (Fig. 3G), and even at the window level (size=100,000, step=20,000), F_ST_ > 0.2 (Fig. 3A). Among these genes, we found *ttc39b, tyrp1, psip1, zdhhc21, frem1,* and *nfib*. This group of genes was initially described to be located in the mouse brown locus (its mutation causes albino phenotypes), which contains the known pigmentation genes *tyrp1* and *bnc2* ^49^. In our *O. pumilio* assembly, the genes are on different scaffolds, but we note that they are also in the same syntenic region of the high-quality *Oophaga sylvatica* genome, suggesting that the synteny is conserved across tetrapods and that these F_ST_ outliers represent the same divergence signal. (Fig. 3G, J).

**Fig. 3.**
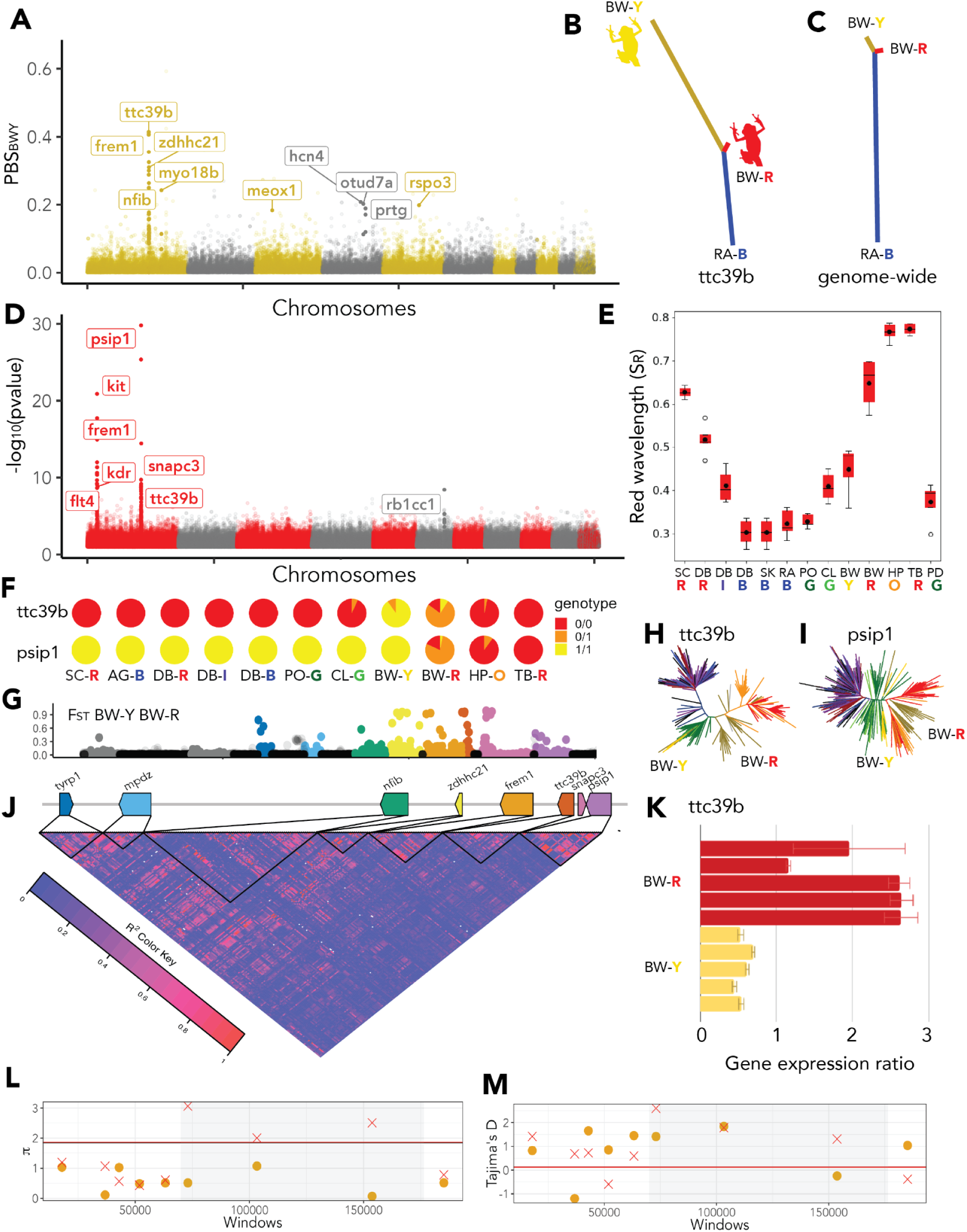
Genetic basis of yellow/red coloration in *O.pumilio.* A) PBS scan of Bastimentos West Yellow branch against BW-R and RA-B B) PBS tree of *ttc39b* scaffold C) PBS genome-wide D) Genomic association to red color wavelengths E) Red wavelength (SR) measurements grouped by population F) Allele frequency of SNP in *ttc39b* is P_RNA_scaffold_1998:131072 located in an intron, Ref C, Alt T, and SNP in *psip1* is P_RNA_scaffold_10726:251470 intronic, Ref T, Alt C. G) Per-site FST of BW-R and BW-Y in the *ttc39b* locus, gene colors correspond to genes in H, gray is other genes and black is intergenic H) *ttc39b* tree, colored according to Fig 1A: BW-Y (yellow), BW-R (gold), CL-G (light green), PP-G (dark green), TB-R (red) I) *psip1* tree J) Linkage disequilibrium map in the *ttc39b* locus K) qPCR results showing gene expression ratio of housekeeping gene *sdha* and *ttc39b* L) Pairwise differences in 1000 sites windows in the *ttc39b* scaffold. The shaded area contains the gene, yellow dots correspond to BW-Y and red crosses to BW-R. The red and black lines represent the genome-wide average in each color morph M) Tajima’s D presented in the same format as panel L.

We used the blue population RA-B as an outgroup in a PBS scan between the Bastimentos West populations (BW-R, BW-Y) to identify which was experiencing allele frequency changes. We found a sharp peak in the BW-Y PBS branch (Fig. 3A, B). *Ttc39b* is the gene with the highest F_ST_ between yellow and red frogs of the polymorphic Bastimentos West population. This gene is involved in carotenoid-based phenotypes in fish and birds ^50–53^. It was also found as a candidate gene with differential expression between different color morphs of other dendrobatids ^54^. The carotenoid that vertebrates obtain through diet is yellow, and red can subsequently be produced by converting the yellow carotenoids into red ketocarotenoids. In birds, it was found that *ttc39b* enhances this ketocarotenoid production ^29^, and our results suggest this mechanism is conserved in frogs.

We performed genomic association tests involving all populations with spectrometric measurements of red wavelength. The smallest p-values were found in *psip1, snapc3,* and *ttc39b* (Fig. 3D, E). Examining the allele frequencies of the SNP with the lowest p-value in *ttc39b*, we found a genotype fixed in most populations, only segregating in Colon (green), Hospital Point (orange), and Bastimentos West populations (yellow/red). All the yellow morphs of Bastimentos West have the alternative allele (Fig. 3F). Given that red is the ancestral color in *O. pumilio*, that only non-red frogs have the alternative, and that we know this gene is involved in yellow-to-red carotenoid production ^29^, this gene’s function is most likely impaired in non-red individuals. We could not detect highly differentiated non-synonymous mutations but found several SNPs in introns with very different allele frequencies between color morphs (Fig. 3F). The average genomic tree does not separate yellow from red Bastimentos West individuals, but the *ttc39b* gene tree separates yellow individuals which, interestingly, cluster within all the green frogs from Colon and Popa (Fig. 3H).

Previous studies using wild pedigrees have found that the color differences in the Bastimentos West population are largely influenced by a single locus with complete dominance of red over yellow ^14^. Our findings strongly suggest that this locus is *ttc39b*. To assess the selection acting on this locus in the Bastimentos West population, we compared π and Tajima’s D against genome-wide levels. There is low π in the yellow morph and increased Tajima’s D (Fig. 3L and M). This pattern is rarely observed but implies very few SNPs segregating at high minor allele frequency. A possible explanation is a recent selective sweep of an admixed haplotype where increased Tajima’s D may be observed in the region around the selected mutation ^55^. An HKA test also supports the hypothesis of selection within the yellow morph (Sup Table 4) with very few polymorphic sites within the BW-Y population (q-value < 0.05). We previously mentioned that the BW-Y *ttc39b* haplotype clusters within the green populations (CL-G, PO-G), which likely share the impairment of red pigment production, leading to green coloration in the presence of blue structural coloring. In contrast, BW-R clusters with red and orange populations HP-O and TB-R (Fig. 3H). A likely explanation for this pattern is that the yellow *ttc39b* haplotype was transferred by admixture from CL-G or PO-G to the Bastimentos West area and then subsequently increased in frequency due to positive selection. A transfer of the *ttc39b* haplotype from the green to the yellow population is also supported by the fact that the yellow haplotype clusters within the green clade in the *ttc39b* tree, while the green clade itself maintains the position observed in the genome-wide tree (see Fig. 3H and Sup Fig. 1).

We did not find any synonymous mutations in *ttc39b*, the mutations with highest F_ST_ are not located in exons. However we found differential expression between BW morphs, with red frogs expressing the gene at a higher level (Fig 3K), suggesting that the differences in *ttc39b* are regulatory.

#### Green

Finding the genes underlying green coloration was more challenging since there is not a green polymorphic population. We carried out genomic association tests with spectrometric measurements of green wavelength (S_G_) and again found *ttc39b* in the lead association peak (Fig. 4A), although SNPs near *psip1* had smaller p-values. The gene trees for *ttc39b* and *psip1* (Fig. 3H, I), suggest that yellow and green frogs share similar alleles in these genes. We also note that green frogs from both Popa and Colon have yellow venters (Sup Fig. 3). This confirms the hypothesis that one aspect of green coloration is yellow pigmentation based on a haplotype shared with the yellow BW-Y population. While we cannot, based on these data, exclude that *psip1* plays a causative role, previous research on the role of *ttc39b* in carotenoid conversion and our expression results (Fig. 3K), suggest a major role for *ttc39b*.

**Fig. 4.**
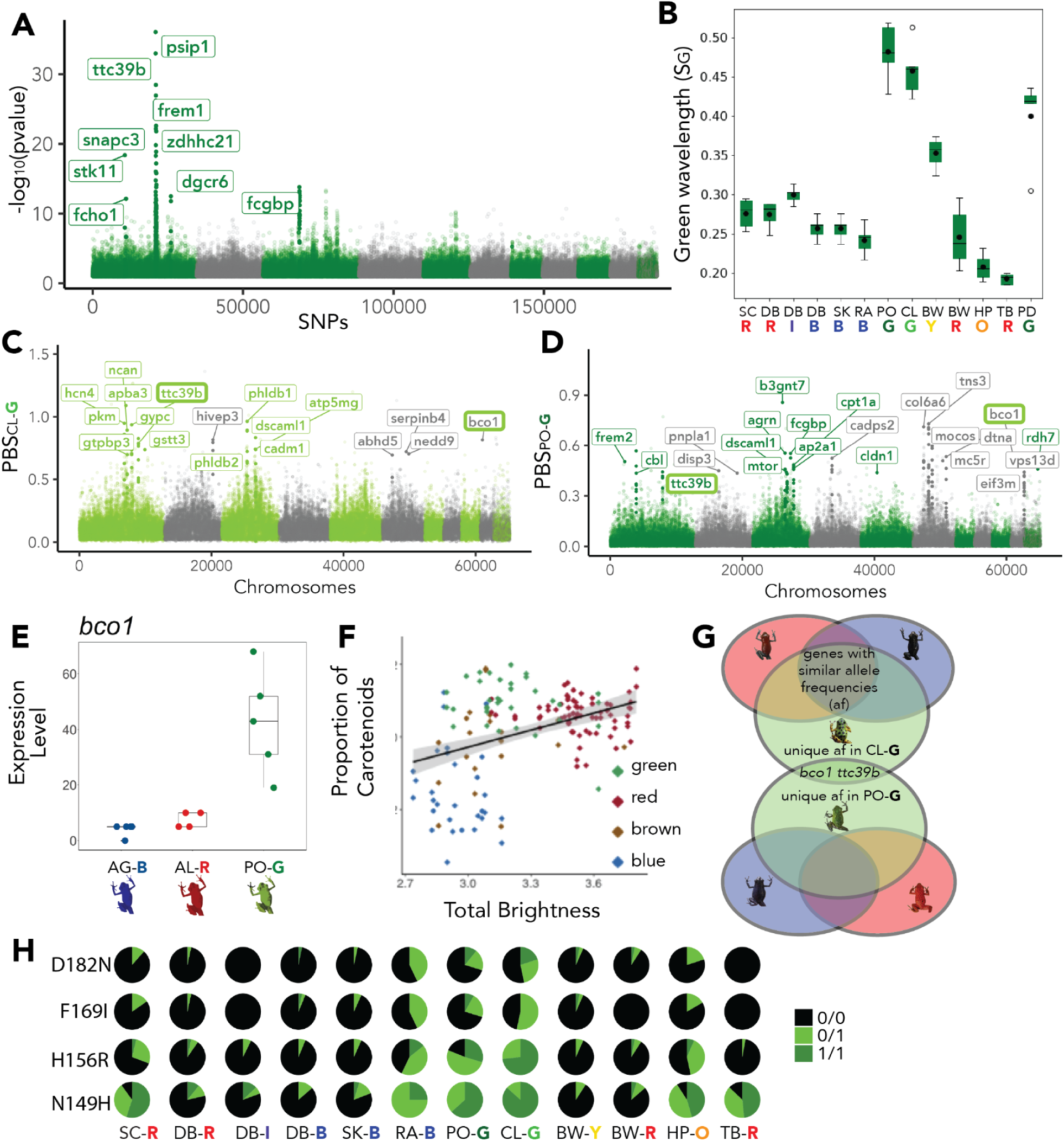
Genetic basis of green coloration in *O.pumilio.* A) Genomic association for green color wavelength (SG) B) Green wavelength measurement by population C) PBS scan of Colon Green branch against DB-B and DB-R D) PBS scan of Popa Green branch against SK-B and TB-R E) Differential expression of *bco1* color morphs F) Proportion of carotenoids in different color morphs of *O. pumilio,* reproduced from Freeborn (2020) G) Venn diagram of the strategy for finding genes differentiated in green frogs H) Non-synonymous mutations found in *bco1*

We conducted PBS scans focused on the branches of the two independent green frog populations, CL-G and PO-G comparing them to non-green populations in two analyses: 1) a PBS analysis of CL-G, versus DB-B and DB-R (Fig. 4C) and 2) of PO-G, versus SK-B and TB-R (Fig. 4D). Besides *ttc39b*, the only other gene we found as an outlier in both scans was *bco1* (Fig. 4G). The expression of *bco1* is much higher in green frogs (Fig. 4E) than in red and blue morphs. *Bco1* is *beta-carotene oxygenase 1,* a gene that cleaves beta-carotenoids into smaller compounds, including apocarotenoids ^56,57^. We also found four non-synonymous mutations segregating in *bco1* with the alternative alleles having a higher frequency in green frogs (Fig. 4H). We found that morphs of *O. pumilio* have different concentrations of beta-carotene in green and red frogs, and apocarotenoid content significantly differs between green and blue morphs (Sup Table 5). Apocarotenoids have a positive linear relationship with spectrometric measurements of green wavelength, and beta-carotenes have a positive linear relationship with red wavelength and brightness (Sup Table 6). Green *O.pumilio* skins have a higher content of carotenoids than blue frogs, and similar content to red frogs (Fig. 4F), but different types (Sup Table 5). Carotenoid processing might be the key to explaining these color differences, making *bco1* a great candidate gene for green coloration.

Other interesting outlier genes in either green PBS scan are *frem2, mc5r*, *fcgbp*, and *col6a6* (Fig. 4C, D). *Mc5r* is one of the five genes in the family of melanocortin receptors in most vertebrates ^58^. This gene is expressed in melanophores and xanthophores and has a role in pigment dispersion ^59–62^. *Mc5r* is differentially expressed in different colors of *O. pumilio* (Sup Fig. 4). *Frem2* has been implicated in melanophore development ^63^. *Fcgbp* has not been functionally found in pigment pathways but was one of the few genes identified to show strong evidence of selection in polar bears compared to brown bears ^64^. We found this gene associated with green wavelength (Fig. 4A). Collagen gene *col6a6* was found as an outlier in multiple PBS analyses: in the green branch of PO-G, SK-B, and TB-R (Fig. 4D), in the blue branch of SK-B, PO-G, and TB-R (Sup Fig. 5), and in the orange branch of HP-O, CL-G, PO-G (Sup Fig. 5). Collagen is important for skin color and its reflective properties in frogs ^40^. The thickness of the collagen layer differs between different colors of *O.pumilio*; red frogs have significantly higher collagen layers than blue frogs ^40^. Thus, our finding of a collagen gene such as *col6a6* in many different comparisons between different morphs could indicate that the gene plays a role in color variation through changes in collagen layer thickness.

#### Pattern

The lead gene in the second highest peak in the association with green wavelength (S_G_) was *stk11* (Fig. 4A). A mutation in this gene causes Peutz-Jeghers syndrome in humans, one of the characteristics of which is pigmentation in the form of freckling or hyperpigmented macules in lips, buccal mucosa, vulva, toes, and fingers ^65–67^. Interestingly, but perhaps coincidentally, the population with the highest S_G_ value is PO-G (Fig. 4B), in which most individuals have a pattern with small speckles, similar to the freckling caused in humans by *stk11* variants. This observation prompted us to investigate the genetic basis of pattern in *Oophaga pumilio*. We did pixel thresholding to convert photographs of frogs into binary format (Fig. 5B), where white pixels correspond to the background dorsal color and black pixels to the pattern (See Methods). We used the proportion of black pixels as a phenotype (BP = black proportion) for a genomic association (Fig. 5A). The most extreme outliers were genes in the *ttc39b* locus. We found that within the polymorphic Bastimentos West population, yellow frogs (BW-Y) have a higher black proportion than red frogs (Fig. 5F). Next, we used a pre-trained convolutional neural network (CNN) for feature extraction on images of frogs (black/white pixels only). We reduced the dimensionality using PCA (Fig. 5D). PC1 distinguishes frogs with small speckles from those with coarse spots (size of spots). PC2, from bottom to top in Fig. 5D, correlates with a gradient ranging from morphs having no pattern to a gradually increasing number of spots (spot density). We did a genomic association with PC1 and found *tyrp1, brms1,* and *rdh8* among the outliers (Fig. 5C). *Brms1* is a metastasis suppressor ^68^ which was also found as an outlier in the BP association. Retinol dehydrogenase (*rdh8*) is a gene involved in visual pigments and retinoid metabolism ^69^. In the association test with PC2, we found *notch1* and *xdh*. *Notch1* was found differentially expressed in *Dendrobates auratus* color morphs and proposed as a good candidate for pattern development in these dendrobatids ^70^. The xanthine hydrogenase (*xdh*), initially described as the rosy locus in *Drosophila*, is known to influence eye coloration ^71^. It is also a gene involved in the pteridine synthesis pathway in amphibians ^27^ that has been shown to be under selection in dendrobatids ^72^. It is also differentially expressed in different color morphs of *D. auratus* ^70^. and *R. imitator* ^54^. Additional analyses suggest that coloration and patterning are not independent of each other (Fig. 5F-H). This may be due to shared molecular mechanisms, or due to conserved linkage between genes involved in patterning (e.g., *tyrp1*) and coloration (e.g., *ttc39b*). For further discussion, see the Supplementary Text.

**Fig. 5.**
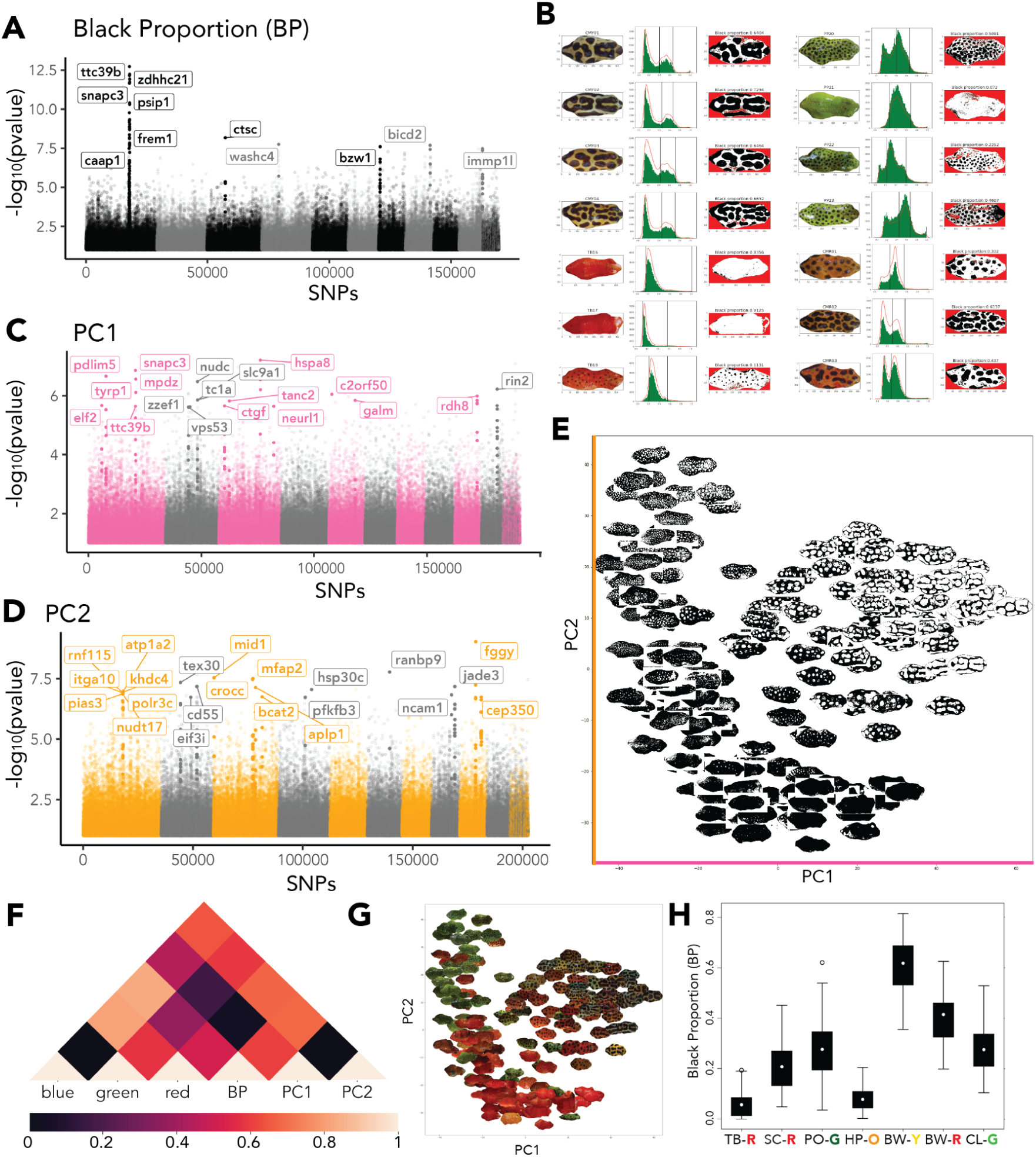
Genetic basis of pattern in *O.pumilio*. A) Genomic association to black proportion. B) Pixel thresholding method examples on different morphs. C) Genomic association to PC1 of feature extraction using VGG16. D) Genomic association to PC2 of feature extraction using VGG16. E) First two components of PCA of VGG16. F) Correlation between pattern (PC1, PC2, BP) and color wavelength measurements (SB, SG, SR). G) Same plot as E but with original colors of images. H) Black proportion in each morph.

## Discussion

In this study, we identified several candidate genes responsible for color and pattern differences in the extremely polymorphic morphs of *O. pumilio* from Bocas del Toro. For blue frogs, the strongest candidate is *kit*. We found two non-synonymous mutations (V533I, G959R) and inferred that the ancestral version of *kit* in this species did not have either of the two mutations, which are also not found in the brighter-colored frogs (Sup Fig. 3). Using the consensus sequence for the gene in each population (Sup Fig. 3), we found that V533I is present in darker populations. Additionally, blue frogs have G959R. The equivalent mutations in humans are associated with cancer (Sup Table 7), as are most described mutations in this gene. More interestingly, a mutation, N975H, makes the C-terminal domain more amphipathic resulting in piebaldism (white patchiness where melanin is missing) in humans^73^. The *O. pumilio* mutation G959R in the C-terminal domain increases hydrophobicity, potentially having the opposite effect, increasing *kit* activation and melanization. This hypothesis is compatible with our finding that blue frogs have more melanophores (Fig. 2G). A sweep in this gene and association with both blue and red wavelengths support the hypothesis that this gene is a key gene that differentiates red frogs from blue.

A single locus dominant for red was previously hypothesized, using pedigree reconstruction from microsatellites, to be responsible for yellow versus red frogs in the Bastimentos West population^14^. Our analyses suggest that this locus includes *ttc39b*, a gene necessary for yellow-to-red carotenoid conversion^29^, which has a shared haplotype between yellow and green frogs. This haplotype might have been introduced by migration from Colon Island into Bastimentos Island. We show strong recent selection occurring in Bastimentos with very few polymorphisms within the yellow frogs (Sup Table 4).

Another important gene with several non-synonymous mutations and higher expression in the green morphs is *bco1*. We hypothesize that this gene is changing the carotenoid content (both quantity and types of carotenoids) in green frogs compared to other colors. By quantifying the different cell types and the amount of blue/green pigments, we provide evidence that both blue and green morphs in *O. pumilio* are produced, at least partly, through structural coloration. Blue frogs are produced through a change in wavelength via an increased number of melanophores (Fig. 2G) and green frogs are produced via different content of carotenoids (Sup Table 5,6). However, detailed functional studies would be needed to confirm the mechanism of action of the associated genes identified in this study.

Many hypotheses involving both selective and neutral evolutionary processes have been proposed regarding the origin and maintenance of color variation in *O. pumilio* in Bocas del Toro ^15,17,74^. We find strong selection signatures in some of our candidate genes. In particular, we find a strong signal of recent selection on a recently introduced haplotype of the *ttc39b* associated with yellow color in the polymorphic population from Bastimentos. This demonstrates that novel phenotypes can be selected for *in situ* within islands after the formation of the islands. We find similar evidence of selection associated with the blue phenotype in the polymorphic Dolphin Bay region of mainland Panama. This suggests that the evolution of new morphs is not necessarily associated with the formation of the islands but instead can be driven by local selection. It also raises two new questions. First, what is the source of this selection? And second, why is its influence limited to the Bocas del Toro area?

While our current study cannot identify the source of selection on color, previous results suggest that avoidance of male-male competition, female mate choice, or viability selection are possible explanations ^6,7,13,20,75^. Male-male competition has recently been shown to be stronger between *O. pumilio* males of the same color morph^6,7,13,20,75^, providing a likely mechanism for selection to favor rare morphs. Why populations throughout the rest of the species range, which includes the Caribbean lowlands of Costa Rica and Nicaragua, are more uniformly red remains a more open question. A possible answer is that the beneficial effects of divergent coloration are transient and that rare morphs eventually go extinct in the larger population on the mainland but may locally fix on islands with smaller population size. There are likely fitness trade-offs between the effects of color on predation, female choice, and male-male competition, and environmental changes may shift the balance between these factors. The Bastimentos West area and Dolphin Bay, where we find evidence of ongoing selection favoring a new morph, are in fact both areas that have been disturbed by humans but yet maintain a very high density of *O. pumilio.* Shifting balance between trade-offs may at times favor rare morphs and at other times favor common morphs, and with different morphs on neighboring islands as reservoirs of genetic variation, a dynamic system might arise that produces the high degree of color variation observed for *O. pumilio* in the Bocas del Toro archipelago. While our results do not provide direct evidence for such dynamics, previous results on male-male competition ^13,20^ and predation ^74^, and our new results demonstrating selection for locally novel color morphs introduced by gene-flow from neighboring islands, support this hypothesis.

Another hypothesis to explain why color variation and polymorphism in *O. pumilio* appears to be limited to the Bocas del Toro area of Panama could be the existence of a locus affecting behavior, where an allele that affects the development of color-dependent male-male aggression and female choice is limited to the Bocas del Toro lineage ^6^. In this case, the selection we observed favoring rare coloration should be limited to lineages that also bear such an “imprinting” allele. More behavioral studies in *O. pumilio*, both within and outside Bocas del Toro, will be needed to investigate this hypothesis. In either case, our results demonstrate that the polymorphisms in Bocas del Toro are driven by natural selection that, at least temporarily, favors rare morphs and cannot be explained simply as a consequence of drift associated with bottleneck during colonization or island formation.

## Acknowledgments

Adam Chois for help cropping frog pictures. Professor Lauren O’Connell for sharing the *Oophaga sylvatica* genome. Ariel Rodriguez for sharing the annotation from their *O. pumilio* rescaffolded genome. NSF DEB 1655336, Collaborative Research: Genomic and phenotypic analyses of color pattern divergence in a mimetic radiation of poison frogs. Fulbright Garcia-Robles and UC MEXUS-CONACYT for supporting DAG while working on this project.

## Data availability

NCBI Database BioProject: PRJNA760522 : *Oophaga pumilio* exomes from Bocas del Toro. All code used in this study is available at https://github.com/aguilar-gomez/pumilioAnalysis.

## Supplementary Materials

Supplemental Text

Materials and Methods

Figures S1 to S7

Tables S1 to S7

Supplementary References

## Supplemental Text

### Additional coloration candidate genes

We found other genes previously related to pigmentation as outliers in different PBS scans. When comparing green to orange frogs, we found a long branch in HP-O for the gene *scarb1* (Sup Fig. 5) ^1–3^. Several keratin genes (*krt10, krt12, krt14, krt17, krt24*) were found as outliers in TB-R ^4^ (Sup Fig. 5). We also found *gnpat, adamts8* and *lyst* as outliers in the PBS scans from RA-B compared to Bastimentos West populations (Sup Fig. 5). *Gnpat* has an adaptive variant in humans that promotes melanin synthesis in Tibetans as an adaptive mechanism against UV radiation, making their skin darker ^5^. *Lyst* has previously been identified as a gene causing lighter skin or coat color phenotypes across many vertebrates, ranging from mammals to reptiles ^6–9^. *Adamts8* has not directly been implicated in pigment phenotypes but has several paralogs involved in chromatophore differentiation and migration, such as *adamts9* ^10^, *adamts13* ^11^, and *adamts20* ^12^ ^13^.

### Additional pattern candidate genes

Some of these genes found in the genetic associations, like *xdh* and *rdh8*, are functionally more related to coloration than patterning. We observed that the black pattern (BP) varied depending on the morph, with some colors and populations having a lot more BP (Figure 6F). Very bright-colored populations such as TB-R and HP-O have very low BP, consistent with previous findings ^14^. We calculated the correlation between all the phenotypes we used for genomic association (color wavelength [blue, green, red], BP, PC1, and PC2) (Figure 6G). PC1 correlates highly with BP, which makes sense since both phenotypes measure patterning. PC2 is also correlated with pattern and additionally to the color wavelengths. Based on these results, we plotted the pictures with color in the PC1 and PC2 axes (Figure 6H). Feature extraction and PCA analysis were performed on binary images (black and white). Therefore, we did not expect that the PCs would separate frogs by color. We can observe how PC2 separates green from orange/red frogs—explaining why we get genes like *xdh* in the association results. It also reinforces that pattern and color are not independent in *O.pumilio* morphs.

*Tyrp1* was found as a candidate for patterning in two associations. As mentioned above, it has already been implicated in patterning phenotypes. Particularly in red tilapia, where red meat is deemed more precious, the phenotype is unstable and often accompanied by black dots, decreasing its commercial value ^2^. In red tilapia, *ttc39b* was found as one of the candidate genes involved in carotenoid storage and *tyrp1* as one of the main genes responsible for the black dots. However, they failed to realize that both genes are in the same locus. The synteny is likely preserved across all vertebrates since we know it is in the same locus in mice and frogs. Another study in common carp patterning found a recent *tyrp1* duplication responsible for patterning. In this system, the fish sometimes have an orange background that is linked to pattern ^15^. *Ttc39b* and *typr1* are likely contributing to all these phenotypes where color and pattern are not independent.

## Materials and Methods

### Sampling

A total of 347 *Oophaga pumilio* frogs were sampled from the Bocas Del Toro Province in Panama. Aguacate blue morph frogs were sampled from the Aguacate Peninsula ( [AG] n=37: Shark Hole n=30 [SK] and Rana Azul n=7l [RA]), which is still part of the mainland but located a short distance from the archipelago. In the same peninsula, we sampled frogs from the blue-red transition zone Dolphin Bay ([DB] n: red=31, blue=32 and intermediate=41). The orange frogs were sampled from Solarte ([HP] n=30, Hospital Point). We sampled frogs from the San Cristobal morph, red with blue legs and sometimes black speckles ([SC] n=26, San Cristobal and South San Cristobal). Popa green frogs were sampled from two locations on Popa Island ([PP] n=37, Punta Laurel [PO] and Popa Dos [PD]). Red frogs were sampled from Bastimentos island in Tranquilo Bay ([TB] n = 35). At the northwestern tip of this same island, we sampled a polymorphic population Bastimentos West (BW). We sampled yellow frogs (BW-Y, n=30) and red frogs (BW-R, n=33) from this area. We also collected green frogs from two locations on Colon Island ([CL] n=15, Boca del Drago and La Gruta). The exact locations are omitted because of the persisting illegal pet trade of *O.pumilio*.

### Probe design for exome capture

The initial transcriptome was 108,640,165 bp represented by 152,862 transcripts. The transcripts were mapped to *the Oophaga pumilio* and the *Ranitomeya imitator* genomes using STARlong 2.7.0d. We removed the transcripts that did not map to these two assemblies. This resulted in 141,482 transcripts and 104,671,021 bp. Then we used transcript_filter.pl v0.2.0 to remove isoforms. We did two rounds of filtering using the following parameters: 1) minimum 80% coverage and 98% identical 2) 90% coverage, 95% identical. The isoform removal step resulted in 97 Mb. We removed most mitochondrial genes using the annotation and kept cytB. We used 6-Process Annotation (https://github.com/CGRL-QB3-UCBerkeley/MarkerDevelopmentPopGen) to filter out sequences smaller than 150 bp and with a GC content smaller than 35% or larger than 75%. This script also masked repeats setting the parameter to “Xenopus genus” and “vertebrates”. Finally, we checked if a list of candidate color genes was present. We added the missing genes from the *Oophaga pumilio* or the *Ranitomeya imitator* genomes. The final design had 90 Mb and 115,420 regions and was sent to NimbleGene for approval. NimbleGene made 116,121 probes that targeted 80 Mb, estimating a final coverage of 86 Mb across 110,329 transcripts.

### Library prep

High molecular weight DNA samples from 347 *Oophaga pumilio* individuals were diluted to obtain volumes between 100 and 170 µL with concentrations lower or equal to 30 ng/µL. Samples with similar concentrations and the same volume were grouped in sonication batches. We sonicated using Q800R3 Sonicator from QSonica. We sonicated DNA samples setting four minutes total ON time, with a pulse of 15 seconds ON / 15 seconds OFF, 40% amplitude. Tubes were spun down in a centrifuge halfway (after two minutes) and restarted. Samples with low concentration (i.e., <1 ng/µL) were sonicated for an extra minute (five minutes total sonication time, same parameters). The target fragment size was ∼300-500bp. We verified that the sonication worked properly by running 1 µL of each sample on a 2% agarose gel.

Sonication was followed by size selection using lab-made SPRI beads [20% PEG-8000 / 2.5M NaCl / 1 mg/mL Sera-Mag SpeedBead Carboxylate-Modified Magnetic Particles (Hydrophobic) 65152105050250 ^16^] to refine the fragment sizing. These magnetic beads reversibly bind to larger or smaller sized DNA fragments depending on the volume added. The target for our libraries was ∼350 bp, so we used a 0.5x ratio for the right-side selection and 0.65x for the left-side. We used the Rainin Benchsmart 96, a high-throughput pipetting system, to treat 96 samples simultaneously. Each plate had randomly selected samples from different populations. An agarose gel was run after the double-sided bead cleaning to confirm the fragment sizes. Then, we used a Kapa HyperPrep library prep kit for end repair and A-tailing, followed by adapter ligation with a universal stub. The next step was a post-ligation bead clean-up to remove excess adapters and ligase. Finally, we performed a 0.8x SPRI cleaning on the Benchsmart.

We needed an indexing PCR to recognize the samples after pooling them together for exome capture and sequencing. We used plates provided by the Berkeley Genome Sequencing Facility that contain a pre-mixed unique P5 and P7 indexing oligo for each sample. The Benchsmart was again used to do a final 0.8x bead clean-up to remove excess indexing oligos and dimers. After adding the adapter and index, the target size of fragments is ∼450-650 bp. After completing the libraries, we did quality control by measuring the final concentration using Qubit and running agarose gels and Bioanalyzer DNA 1000 chips to obtain library sizing information.

### Pooling and exome capture

We made five pools containing ∼70 samples each: we pipetted different volumes per sample depending on the concentration to ensure we had the same mass for each sample (150 ng). Each pool was constituted of samples from different populations and contained ∼3 µg of DNA. We added 15 µL of blocking oligos (Roche UBO), 5 µL of Cot-1 from each of three different species (human, mouse, chicken) and 15 µL of Roche Developer Reagent. Then we concentrated the pools in a vacuum centrifuge and resuspended the pools in Roche Nimblegen buffers 5 and 6. We aliquoted the Nimblegen probes into strip tubes, added the DNA mix to the probes, denatured at 95°C, and started hybridizing at 47°C. We incubated for 72 hours and 45 minutes. After probe hybridization, we followed the Nimblegen protocol to bind the proves to streptavidin beads, thoroughly wash the beads, elute the captured DNA, and proceeded to post-capture PCR amplification using 2x Kapa HiFi HotStart ReadyMix for ten cycles. We assessed the final concentration, size distribution, and quality of the amplified captured multiple DNA samples using Qubit values, Bioanalyzer, and agarose gel electrophoresis.

### Sequencing and Read processing

We sequenced two lanes of Illumina NovaSeq 6000 150PE Flow Cell S4. We used *Trimmomatic-0.39* to remove adapters (ILLUMINACLIP:TruSeq3-PE.fa:2:30:10), leading and trailing low quality or N bases (below quality 3) (LEADING:3,TRAILING:3 ), cut reads when the average quality per base drops below 15 in a 4-base sliding window (SLIDINGWINDOW:4:15) and drop reads below the 75 bases long (MINLEN:75). We also used PRINSEQ-lite 0.20.4 to remove low complexity reads (-lc_method dust -lc_threshold 7) and overrepresented sequences (-custom_params “G 50”).

### Mapping

The reads from each sequencing lane were mapped independently to the *Oophaga pumilio* rescaffolded genome [total length: 4,837,165,062 bp/ L50: 11,434/ N50: 105,380 bp] ^1,17^ using bwa mem 0.7.17-r1188 ^18^ . Next, we filtered the alignments with samtools 1.11 ^19^ and used the flag -F 1804 to exclude reads: read unmapped (0x4), mate unmapped (0x8), non-primary alignment (0x100), read fails platform/vendor quality checks (0x200) and read is PCR or optical duplicate (0x400). Finally, we merged the bam files coming from different lanes for each individual.

### SNP filtering

SNP filtering was performed using ANGSD *0.941-13-gb7eb654* ^20^, we kept SNPs with a minimum of 250 individuals with 1x depth (minInd, setMinDepthInd), maximum depth of 50x (setMaxDepthInd), base quality 25 (minQ), mapping quality 25 (minMapQ), polymorphic sites p-value<1e-6 (SNP_pval), remove triallelic sites and use the reference genome as major (skipTriallelic, doMajorMinor), discard reads with flag >=256 (remove_bads), discard reads that do not map uniquely (uniqueOnly), keep only paired-end reads that mapped correctly (only_proper_pairs). We also removed SNPs with excess heterozygotes (hetbias_pval 1e-6), strand bias (sb_pval 1e-4), end distance bias (edge_pval 1e-4), and map quality bias (mapQ_pval 1e-4). This filtering resulted in 10,231,695 sites. These sites have an average depth of 8.22x.

### Population structure analyses

A 5% minor allele frequency was implemented to run the population structure analyses. We obtained 1,973,099 SNPs after this filter. We used *PCAngsd 1.10* ^21^ to perform Principal Component Analysis (PCA). The program *qpas* from *OHANA* ^22^ was used to calculate ancestry components with k=4 to k=10. We used 1% of the sites to calculate ancestry components. We calculated the covariance of the allele frequencies in different components using *nemeco* to approximate a tree.

### Demographic inference

We used a vcf with genotype calls from bcftools with the SNPs filtered as described above. We used a script available on GitHub to convert to TreeMix format (https://github.com/speciationgenomics/scripts/blob/master/vcf2treemix.sh). We ran AdmixtureBayes on the input file for TreeMix when allowing for each of 0, 1, 2, 3, and 4 admixture events, and obtained a distribution of admixture graphs for each number of events. Three independent MCMC chains were run for each analysis, and convergence was assessed through both trace plot analysis (Sup. Fig 6) and Gelman-Rubin convergence diagnostics (Sup. fig 7) to verify that the stationary distribution had been reached. Each chain was run with the parameters --n 750000 --MCMC_chains 31 --spacing 1.4 --maxtemp 1000 --max_admixes k --num_ind_snps 40000, where k denotes the number of admixture events that were allowed. For the *analyzeSamples* step, a burn-in fraction of 0.3 and a thinning rate of 10 were used to obtain our samples of graphs from the stationary distribution.

We calculated the folded 2dsfs and ran *dadi* ^23^ with pairs of populations using two models: 1) population split and 2) population split with migration. The mutation rate was set to 10^-9,^ and the generation time was set to two years ^24^. The total length of DNA sequence analyzed to obtain the SNP data (10,788,793 bp) was set to 164,229,310 bp, which includes invariant sites. This number was obtained from mapping the exome capture data to the whole genome and keeping only positions that passed the following thresholds: 250 individuals with 1x depth (*minInd, setMinDepthInd)*, maximum depth of 50x (setMaxDepthInd), base quality 25 (*minQ*), mapping quality 25 (*minMapQ*). We generated the folded joint SFS for each pair of populations using realSFS. We ran the program 100 times for each pair of populations and selected the model with the highest log-likelihood. We used the split with migration model, which has the following parameters: population sizes (nu1,nu2), time of the split (T), and migration rates (m12, m21, we used symmetric migration). The initial parameters for the models were (params = [1,1,1,1], lower_bounds = [1e-3, 1e-3, 1e-3, 1e-5], upper_bounds = [200,200, 300, 100]). For the model with no migration we used the same parameters and set m12=m21=0. We used the method to perturb the parameters for each run.

### Selection scans and genomic associations

We calculated the population divergence using *realSFS* to estimate the fixation index F_ST_. We performed selection scans by calculating the Population Branch Statistic (PBS) by windows across the genome using *realSFS*. We calculated the unfolded 2dsfs of the three pairs of each combination. We obtained a global estimation of F_ST_ and PBS, followed by window estimation (size=100,000, step=20,000). We annotated the 99.8th outliers with windows with at least 20 SNPs.

Genomic associations were performed on spectrophotometer measurements of color wavelengths. We used GEMMA 0.98.5 to perform a Wald test using a standardized relatedness matrix (calculated with gemma). We filtered out genes with p-value>0.1 for plotting and selecting outliers. We annotated the 99.98th percentile outliers and kept only genes with at least two outlier SNPs. We aligned the *O.pumilio* genome to the contig level *Ranitomeya imitator* ^4^ genome and plotted our scaffolds using that synteny.

### Differential expression

Publically available RNA-seq skin cell dataset from five green, five red, and five blue Bocas del Toro *Oophaga pumilio^1^* was downloaded using fasterq-dump from NCBI.mTrimmomatic (v0.39) parameters: “2:30:10:2:keepBothReads LEADING:3 TRAILING:3 MINLEN:75” was used to remove adapters. We generated a gtf file using gffread (v0.12.7) from the gff annotation without gene names provided by the author of the publicly available dataset. We used STAR’s “--runMode genomeGenerate” function to index the genome using the following parameter “--limitGenomeGenerateRAM 4272279470500 --genomeSAindexNbases 15.086”. Afterwards, we aligned the reads to the indexed reference genome using STAR (v2.7.10): “--runMode alignReads” with parameter “--readFilesCommand zcat --quantMode TranscriptomeSAM”. Transcript quantification was done using RSEM v1.3.3 . with rsem-prepare-reference followed by rsem-calculate-expression.

For the annotation, we used TransDecoder (v5.5.0)’s gff3_file_to_bed.pl to transform the gff file into a bed file. We blasted all the transcripts since the gff was provided without gene names. We kept the gene name annotation for each transcript with the highest bitscore. For our genes of interest, we also corroborated the annotation of the transcripts and the genome assembly location using our assembly annotation. We ran differential expression analysis with DESeq2 in R (v4.2.0). Transcripts were aggregated by gene name. We subset to only candidate genes that were outliers of selection, color association genomic scans and had a previous link to pigmentation (14 of genes). We did all the pairwise combinations (blue-red-green) and ran the DESeq2 analysis function. We kept the significant DE genes (q-value<0.05).

### Image processing

For this project, frog pictures were not taken with the original intent of using them for image analysis. Therefore the quality of some of them could have been more optimal (lightning, sharpness). We excluded blurry pictures. Since the image analysis aimed to understand the pattern and not the color, we increased the contrast of some pictures (Adobe Photoshop), which may sometimes alter the colors. We excluded all the frogs from the Aguacate Peninsula (RA, SK, and DB) since this region’s dark hue and the pictures’ quality would not allow us to analyze them properly. The frogs were cropped using Pixlr E (https://pixlr.com/e/) to remove all the frogs’ backgrounds and limbs. We also removed the eyes if visible in the picture to avoid confusion with the black spotted pattern. We rotated all pictures to have the frog body lay horizontally.

We used the Python library *scikit-image* ^25^ to adjust the exposure using two consecutive logarithmic corrections. We then converted the images to grayscale. Transparent pixels, which by default are assigned a zero value, were converted to “nan” because the end goal was to convert the pictures into binary black (0) and white (1) images. We used thresholding segmentation. Based on the distribution of pixels, the cutoff is selected where the slope of the distribution changes. We use the cutoff to divide the pixels into binary values (0,1) (Figure Pattern B). We also noticed that since our pictures were not taken for image analysis purposes, some of them contain glare, making some parts of the picture white. Our algorithm also includes a second threshold to exclude these bright pixels. We used the proportion of black pixels to calculate each frog’s Black Proportion (BP) value. These values were used as a phenotype for a genomic association.

### Pattern classification

We tried to understand and classify frogs based on patterns. We used the processed images described above, binary images of the cropped frogs with glare removed in format png. Then we did feature extraction using a pre-trained convoluted neural network with 16 layers called VGG-16 ^26^. We then used PCA for dimensionality reduction of the extracted features. All the analysis was performed in python using libraries keras, scikit-learn, numpy, and scikit-image.

### HKA test

We used the HKA test ^27^ to assess whether the genes we found involved in pigmentation have neutral evolution patterns. We calculated the allele frequencies using angsd (-doMaf 1) ^28^ and the allele in the reference genome as major (-doMajorMinor 4). We kept non-variable sites and biallelic variants that passed the above SNP filtering. We compared *O. pumilio* population allele frequencies against the outgroup *R.imitator*. A site was defined as polymorphic when segregating (.001< af1 <.999 ) in the population of interest and fixed (af2 <.001 or af2 >.999) in the outgroup. A site was defined as fixed if the population and the outgroup were fixed for different alleles. By adding a divergent outgroup, some of our biallelic SNPs in *O.pumilio* were triallelic, with a different allele in *R. imitator*. We also counted a site as both fixed and polymorphic if it was segregating in *O.pumilio* and fixed for a third allele in the outgroup. We counted all the fixed and polymorphic sites genome-wide. We built a contingency table comparing the fixed and polymorphic sites within a scaffold compared to genome-wide counts and performed a X^2^ test using Yates correction. We adjusted the X^2^ values using genomic control and calculating a deflation/inflation index.

### Tajima’s D

We calculated site allele frequency likelihood using *doSaf -1* of all the sites that passed the SNP filters and the invariable sites that passed the mapping and read quality filters. Then we calculated the maximum likelihood estimate of the folded site frequency spectrum (realSFS) and thetas estimate for each site using saf2theta ^29^. We calculated Tajima’s D according to the formula described in ^30^. We performed the calculations using windows of 1000 sites.

### Synonymous and Non-synonymous mutations

We generated a consensus fasta sequence per population (-doFasta 2 from ANGSD). Then we used the gff3 file to extract the CDS and compare the sequences between morphs. We obtained the synonymous and non-synonymous mutations in the candidate genes. We explored the structure of the protein changes using SWISS-MODEL (https://swissmodel.expasy.org/).

---

**Sup Fig 1.**
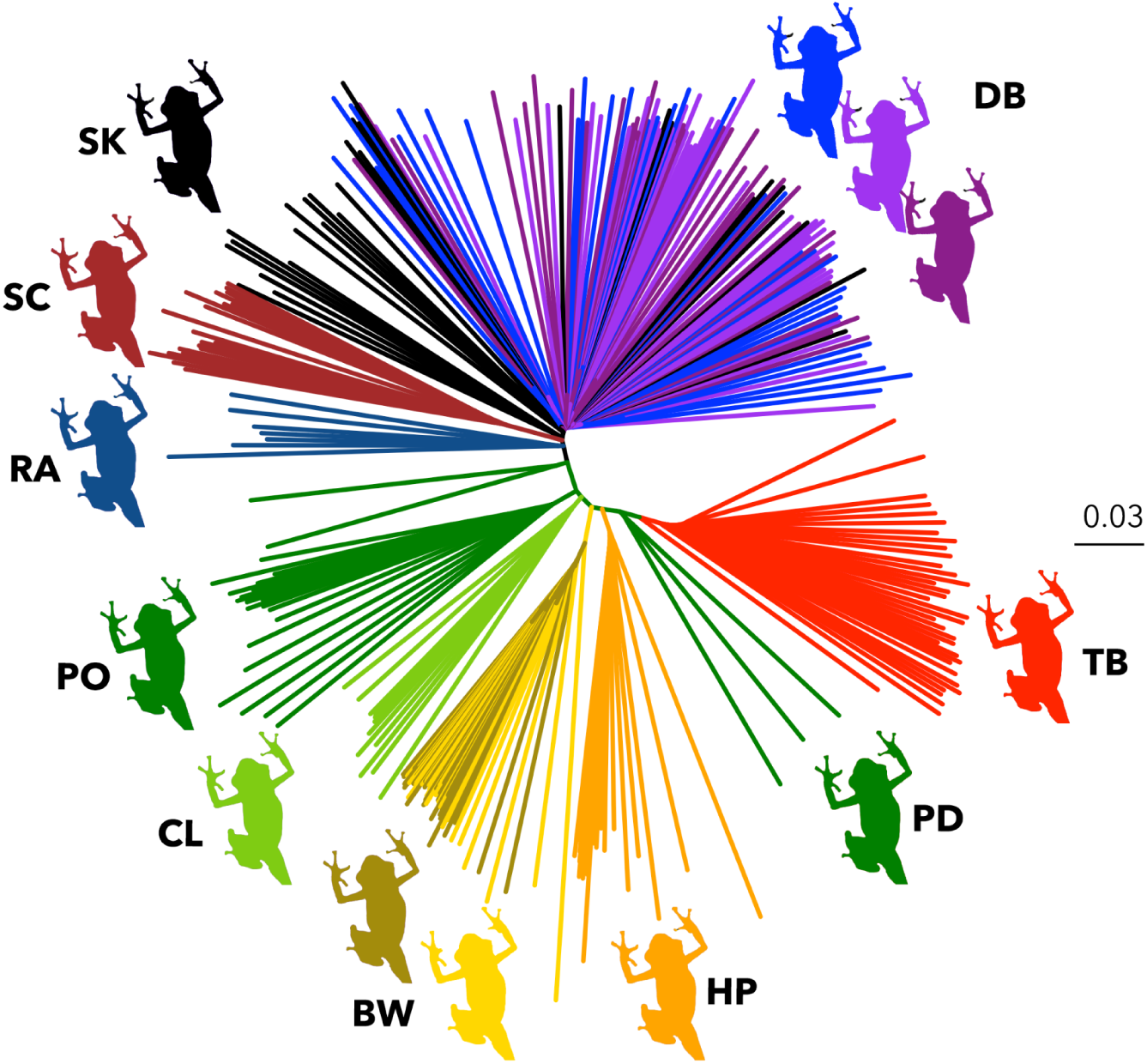
Average Genomic Tree.

**Sup Fig 2.**
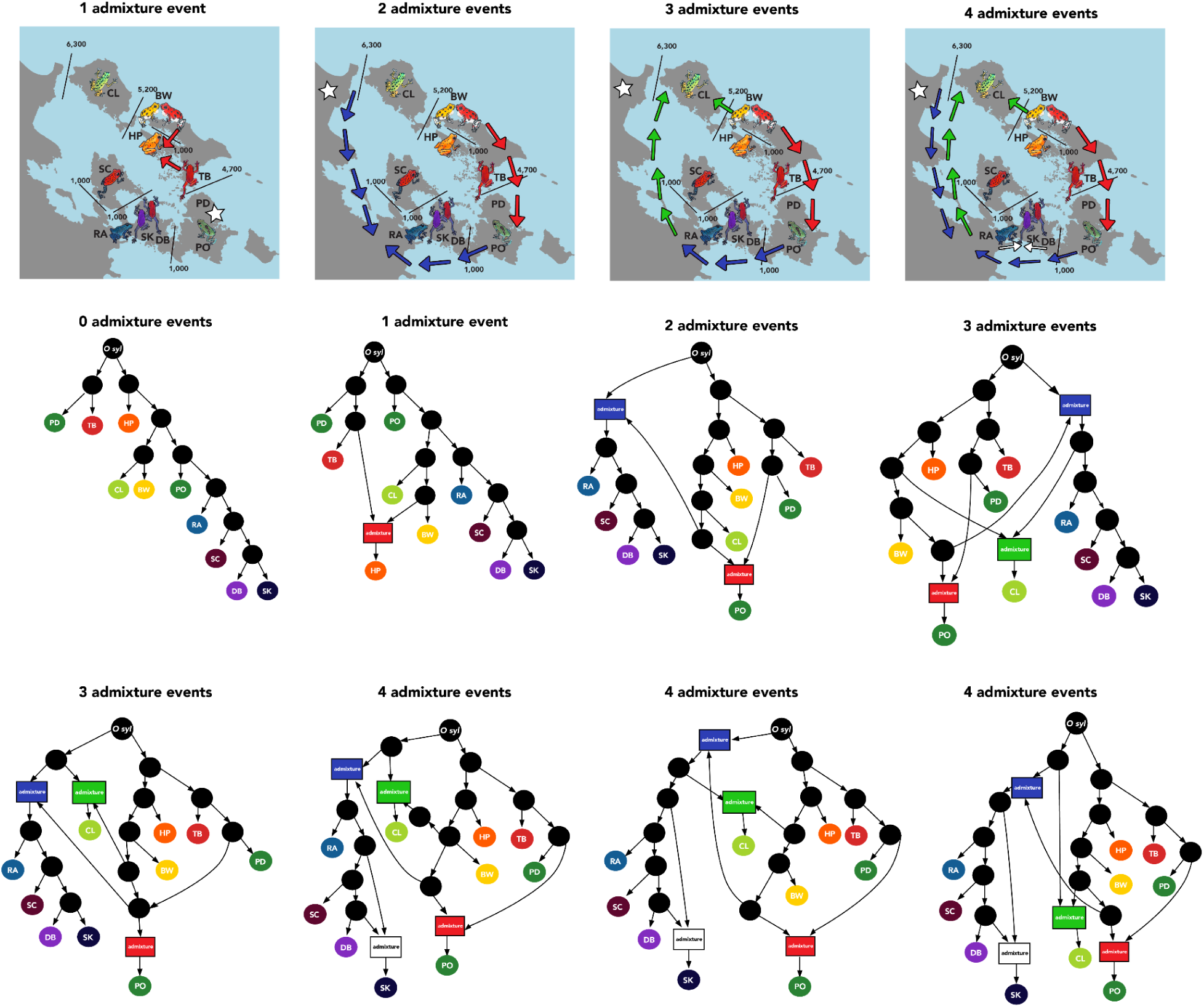
AdmixtureBayes. Topologies and their probabilities given the number of migration events. 0 events (100%), 1 event (100%), 2 events: (100%), 3 events: (68%, 32%) and 4 events (45%, 17%, 12%, the probability of these three topologies is 74% of all sampled graphs with 4 migration events, the rest of the topologies each have 1-3% probability and are not shown here).

**Sup Fig 3.**
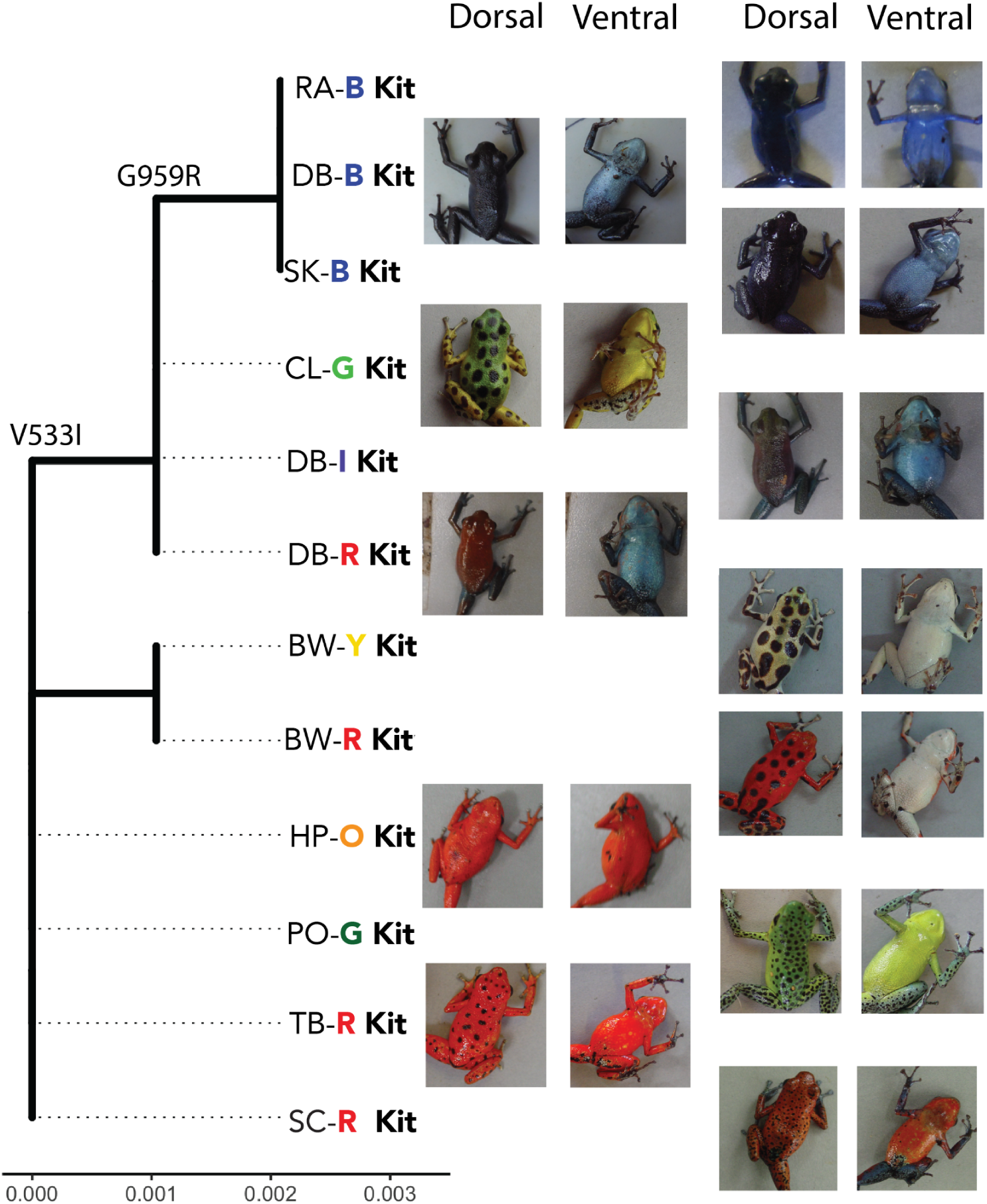
Kit amino acid tree with dorsal and ventral coloration of all populations

**Sup Fig 4.**
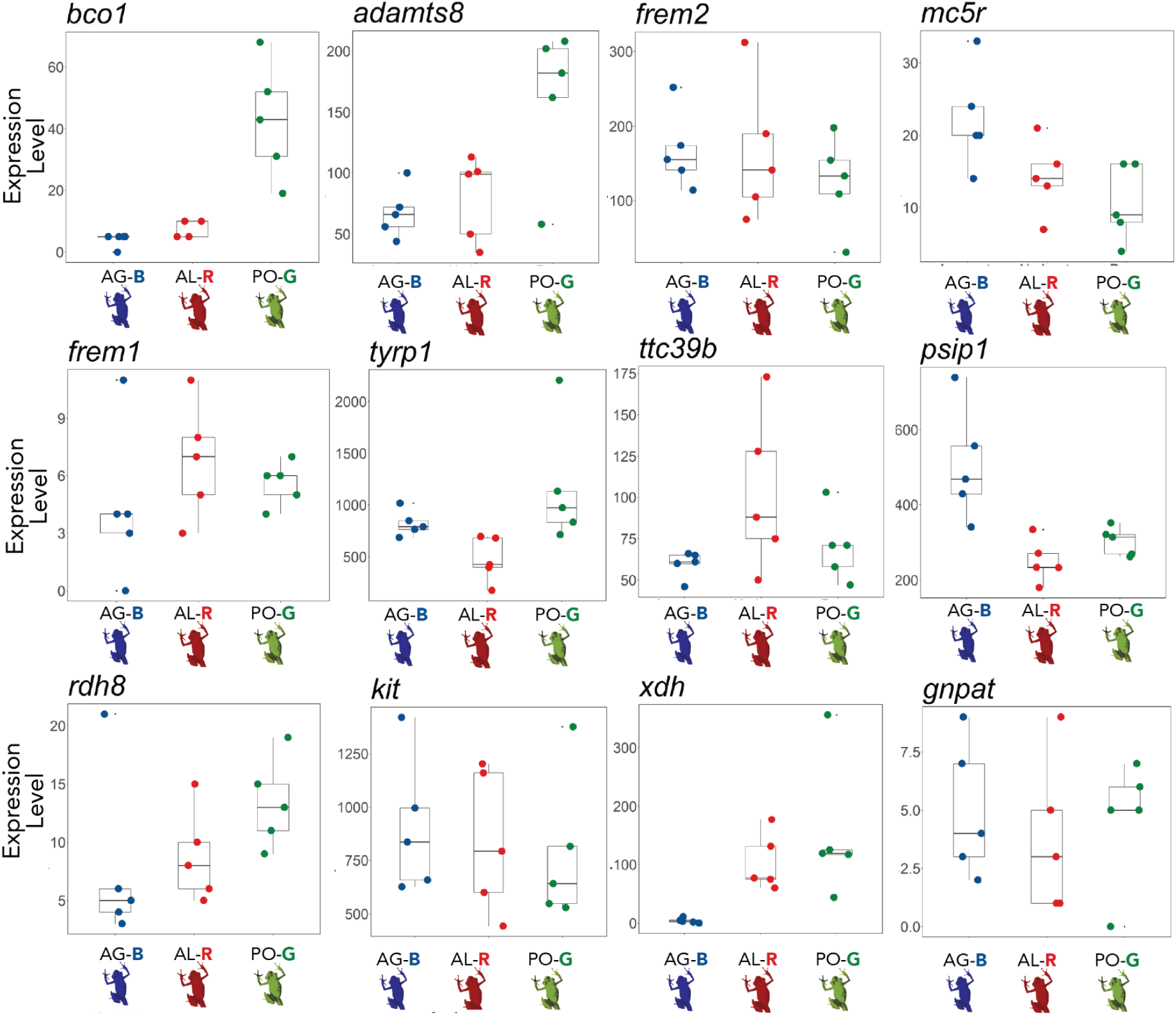
Differential expression of candidate genes

**Sup Fig 5.**
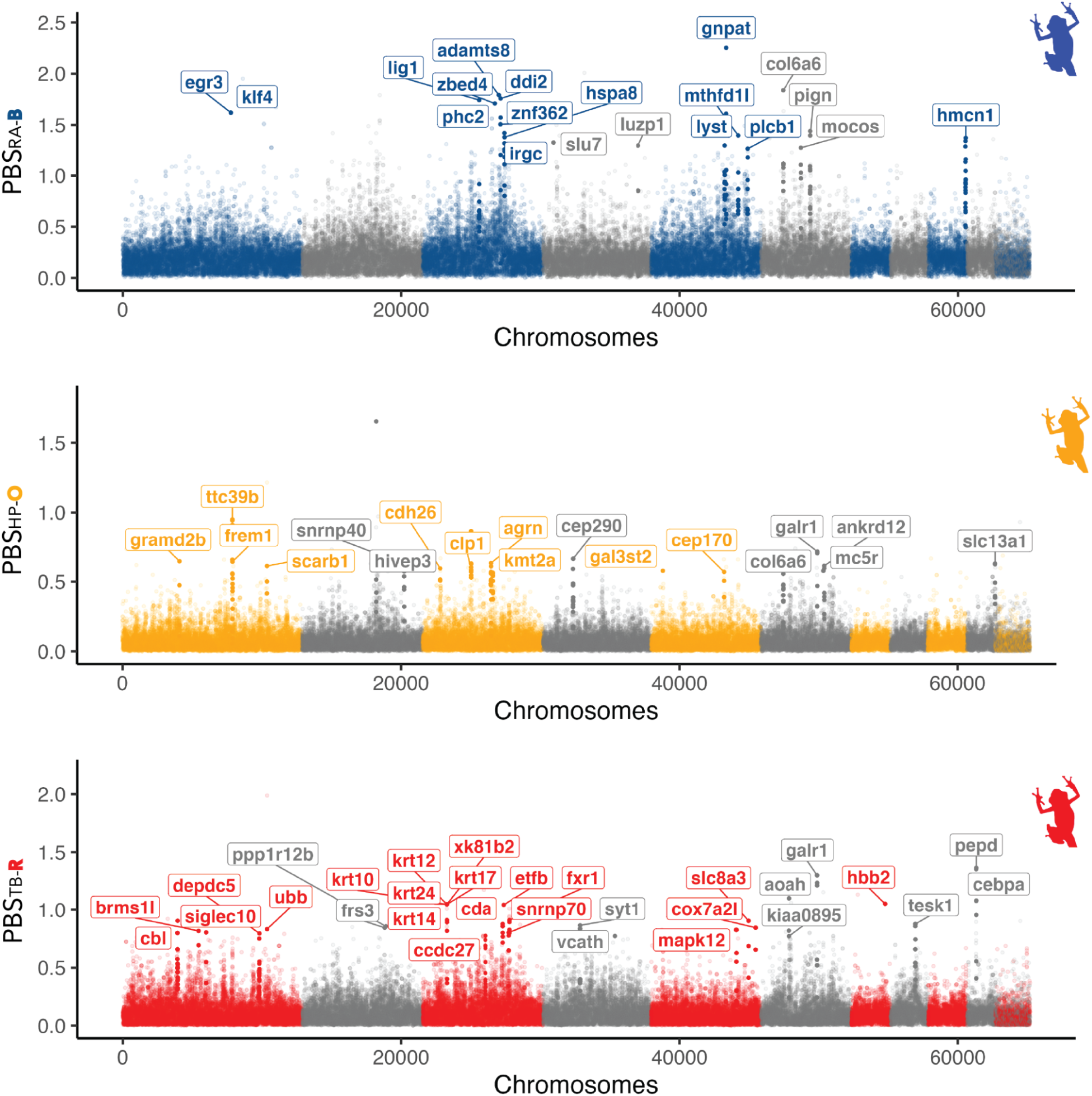
Additional PBS scans. From top to bottom: RA-B vs BW-R and BW-Y, HP-O vs CL-G and PO-G, TB-R vs SK-B and PO-G

**Sup Figure 6.**
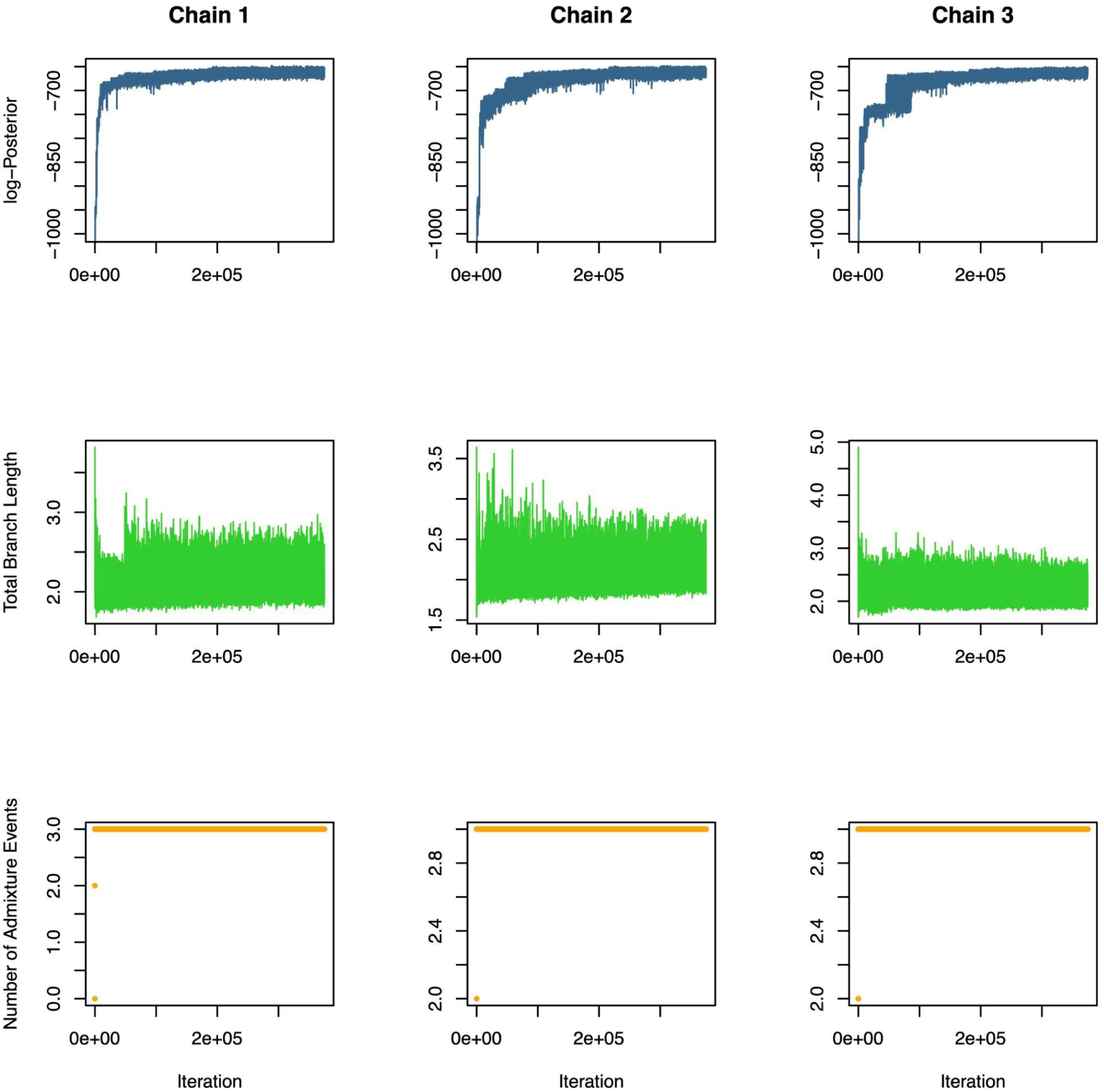
Convergence of Admixture Bayes. Representative trace plots of summary statistics of admixture graphs to assess the convergence of the Markov chain. This figure shows the trace plots for our 3 admixture event analyses.

**Sup Figure 7.**
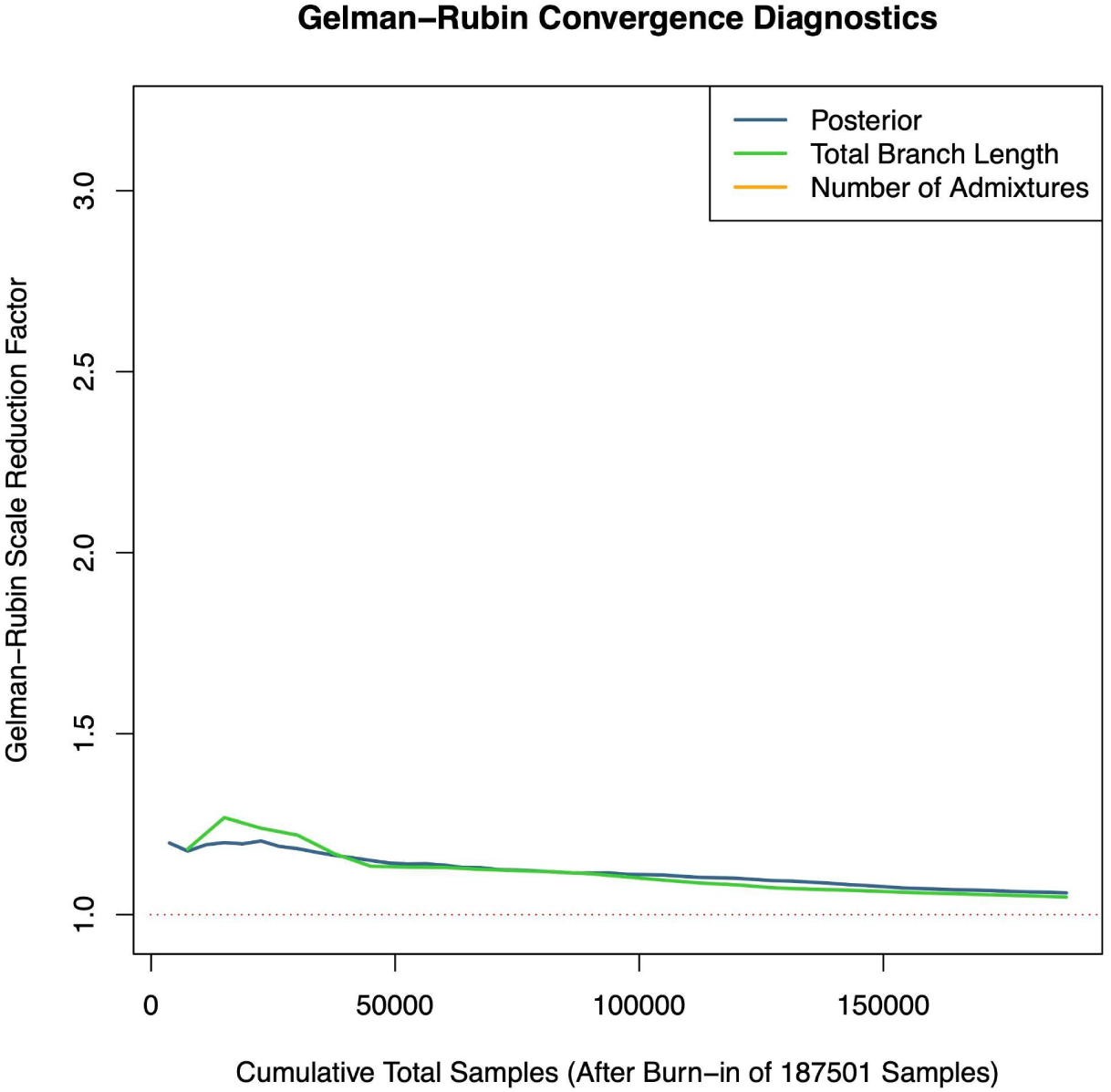
Gelman-Rubin convergence diagnostics. Used to assess the convergence of the Markov chain. This figure shows the Gelman-Rubin statistics for our 3 admixture event analyses. Convergence to 1 indicates that all chains have reached the stationary distribution of the Markov chain

**Sup Table 1.** Pairwise F_ST_ between Bocas del Toro populations and morphs.

|  | SK | RA | CL | BWR | BWY | DBB | DBI | DBR | HP | PD | PO | SC | TB |
| --- | --- | --- | --- | --- | --- | --- | --- | --- | --- | --- | --- | --- | --- |
| SK | - | 0.054 | 0.126 | 0.194 | 0.193 | 0.012 | 0.011 | 0.012 | 0.178 | 0.191 | 0.124 | 0.132 | 0.202 |
| RA | 0.054 | - | 0.137 | 0.207 | 0.208 | 0.06 | 0.058 | 0.06 | 0.177 | 0.193 | 0.125 | 0.146 | 0.187 |
| CL | 0.126 | 0.137 | - | 0.134 | 0.135 | 0.131 | 0.127 | 0.131 | 0.134 | 0.183 | 0.114 | 0.281 | 0.173 |
| BWR | 0.194 | 0.207 | 0.134 | - | 0.009 | 0.199 | 0.188 | 0.202 | 0.152 | 0.232 | 0.156 | 0.281 | 0.21 |
| BWY | 0.193 | 0.208 | 0.135 | 0.009 | - | 0.197 | 0.186 | 0.2 | 0.152 | 0.234 | 0.155 | 0.279 | 0.207 |
| DBB | 0.012 | 0.06 | 0.131 | 0.199 | 0.197 | - | 0.009 | 0.01 | 0.182 | 0.194 | 0.13 | 0.137 | 0.207 |
| DBI | 0.011 | 0.058 | 0.127 | 0.188 | 0.186 | 0.009 | - | 0.008 | 0.178 | 0.192 | 0.128 | 0.135 | 0.206 |
| DBR | 0.012 | 0.06 | 0.131 | 0.202 | 0.2 | 0.01 | 0.008 | - | 0.183 | 0.195 | 0.131 | 0.136 | 0.207 |
| HP | 0.178 | 0.177 | 0.134 | 0.152 | 0.152 | 0.182 | 0.178 | 0.183 | - | 0.159 | 0.142 | 0.253 | 0.154 |
| PD | 0.191 | 0.193 | 0.183 | 0.232 | 0.234 | 0.194 | 0.192 | 0.195 | 0.159 | - | 0.139 | 0.257 | 0.126 |
| PO | 0.124 | 0.125<br>281 | 0.114 | 0.156 | 0.155 | 0.13 | 0.128 | 0.131 | 0.142 | 0.139 | - | 0.222 | 0.164 |
| SC | 0.132 | 0.146 | 0.195 | 0.281 | 0.279 | 0.137 | 0.135 | 0.136 | 0.253 | 0.257 | 0.222 | - | 0.271 |
| TB | 0.202 | 0.187 | 0.173 | 0.21 | 0.207 | 0.207 | 0.206 | 0.207 | 0.154 | 0.126 | 0.164 | 0.271 | - |

**Sup Table 2.** 4-population test .Here are the tests mentioned in the text. The complete results of all the possible combinations for the 4-population test are included in a separate spreadsheet.

| D | JK-D | V(JK-D) | Z | p-value | nABBA | nBABA | nBlocks | H1 | H2 | H3 | H4 |
| --- | --- | --- | --- | --- | --- | --- | --- | --- | --- | --- | --- |
| 0.046 | 0.046 | 3.00E-06 | 26.59 | 0 | 26903 | 24555 | 51422 | HP | TB | PD | *O. sylvatica* |
| -0.009 | -0.009 | 3.00E-06 | -5.37 | 0 | 24093 | 24555 | 53535 | PD | TB | HP | *O. sylvatica* |
| 0.058 | 0.058 | 2.00E-06 | 38.50 | 0 | 22297 | 19851 | 53972 | CL | BW | HP | *O. sylvatica* |
| -0.107 | -0.107 | 3.00E-06 | -66.37 | 0 | 19851 | 24629 | 53350 | BW | HP | CL | *O. sylvatica* |
| 0.060 | 0.060 | 2.00E-06 | 38.76 | 0 | 23340 | 20717 | 52584 | CL | PO | RA | *O. sylvatica* |
| -0.028 | -0.028 | 3.00E-06 | -16.52 | 0 | 20717 | 21906 | 52594 | PO | RA | CL | *O. sylvatica* |
| -0.049 | -0.049 | 3.00E-06 | -26.84 | 0 | 22835 | 25181 | 53558 | PD | PO | TB | *O. sylvatica* |
| -0.035 | -0.035 | 3.00E-06 | -19.04 | 0 | 25181 | 26995 | 51977 | PO | TB | PD | *O. sylvatica* |
| -0.033 | -0.033 | 2.00E-06 | -20.88 | 0 | 21012 | 22430 | 54613 | BW | PO | HP | *O. sylvatica* |
| 0.031 | 0.031 | 3.00E-06 | 18.79 | 0 | 23858 | 22430 | 54551 | HP | PO | BW | *O. sylvatica* |
| -0.025 | -0.025 | 2.00E-06 | -19.34 | 0 | 21779 | 22896 | 53383 | DB | RA | SC | *O. sylvatica* |
| -0.015 | -0.015 | 3.00E-06 | -9.30 | 0 | 22896 | 23583 | 54445 | RA | SC | DB | *O. sylvatica* |

**Sup Table 3.**
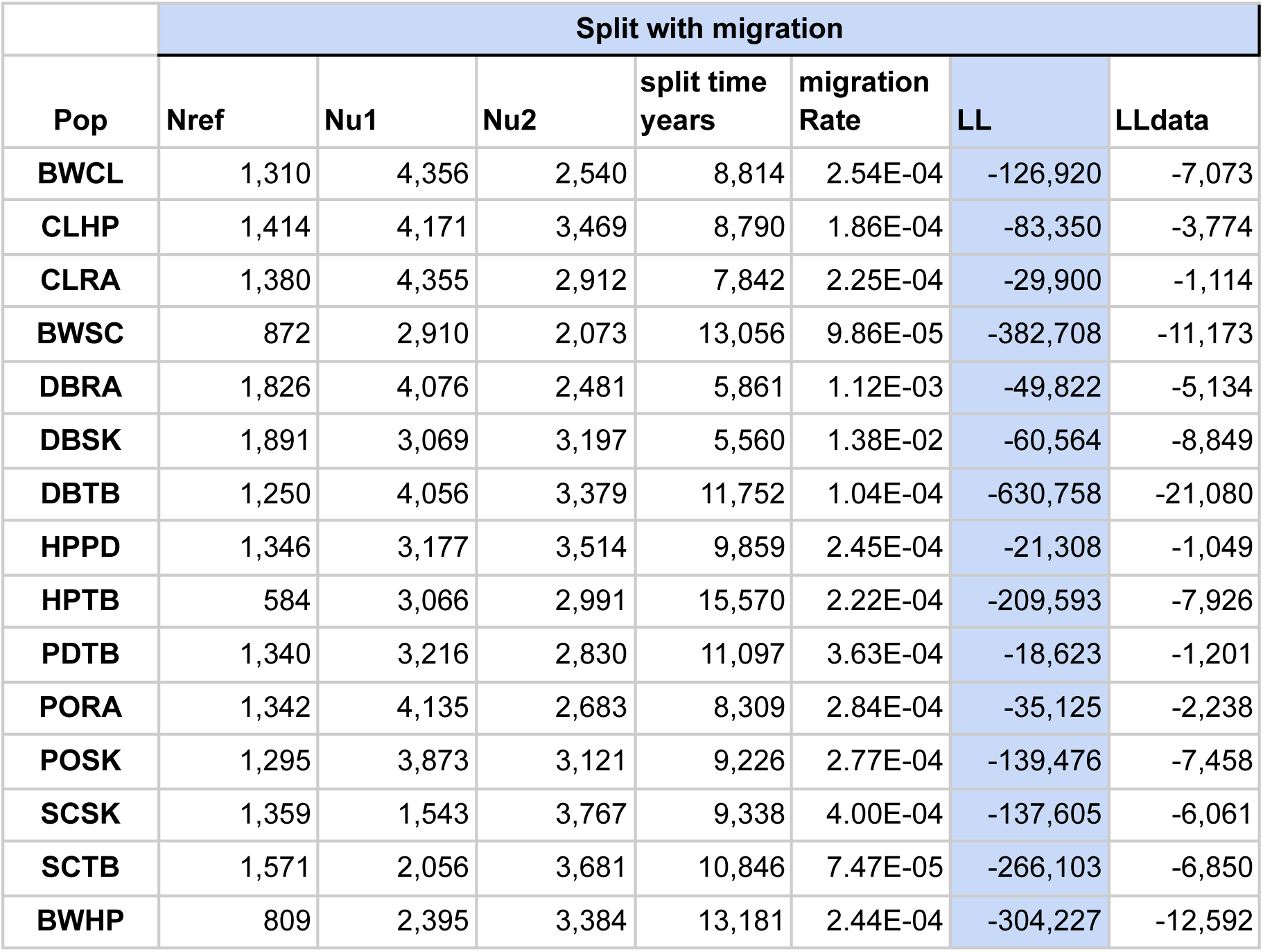
Demographic model results.

**Sup Table 4.** HKA test.

| Population | Color | Pop | gene | Polymorphic gene | Fixed gene | Total sites | corrected chi2 | qvalue | significant |
| --- | --- | --- | --- | --- | --- | --- | --- | --- | --- |
| Bastimentos West | All | BW | ttc39b | 27 | 91 | 118 | 2.4085 | 0.1207 | FALSE |
|  | Red | BW-R |  | 25 | 91 | 116 | 1.5884 | 0.2076 | FALSE |
|  | Yellow | BW-Y |  | 19 | 96 | 115 | 3.8431 | 0.0499 | TRUE |
| Colon | Green | CL-G |  | 17 | 96 | 113 | 4.8832 | 0.0271 | TRUE |
| Bastimentos West | All | BW | tyrp1 | 43 | 130 | 173 | 2.4855 | 0.1149 | FALSE |
|  | Red | BW-R |  | 37 | 130 | 167 | 2.0461 | 0.1526 | FALSE |
|  | Yellow | BW-Y |  | 18 | 138 | 156 | 9.8804 | 0.0017 | TRUE |
| Colon | Green | CL-G |  | 19 | 137 | 156 | 9.5423 | 0.0020 | TRUE |
| Dolphin Bay | All | DB | kit | 105 | 170 | 275 | 0.3854 | 0.5347 | FALSE |
|  | Red | DB-R |  | 76 | 171 | 247 | 0.5117 | 0.4744 | FALSE |
|  | Blue | DB-B |  | 60 | 171 | 231 | 0.3300 | 0.5657 | FALSE |
| Shark Hole | Blue | SK-B |  | 54 | 174 | 228 | 5.9185 | 0.0150 | TRUE |

**Sup Table 5.** Carotenoid differences between *O.pumilio* of different dorsal colors. Reproduced from Freeborn (2020)

| Comparison | Carotenoid | Significance of SIMPER test |
| --- | --- | --- |
| Blue vs green | apocarotenoid | 0 |
|  | canary xanthophyll ester 1 | 0 |
|  | canary xanthophyll ester 2 | 0 |
|  | canary xanthophyll ester 3 | 0.01 |
|  | ketocaritenoid ester 2 | 0.001 |
| Blue vs red | beta-carotene | 0.01 |
|  | canthaxathin ester 1 | 0.01 |
|  | cis lutein ester 1 | 0.01 |
| Green vs Red | beta-carotene | 0.01 |
|  | cis lutein ester 1 | 0.01 |

**Sup Table 6.** Linear relationship between carotenoids and colorimetric variables. Reproduced from Freeborn (2020). Significance codes: 0 ‘***’ 0.001 ‘**’ 0.01 ‘*’ not significant ‘ns’. B: brightness, S_R_: red, S_Y_: yellow, S_G_: green, S_B_: blue, S_V_: violet.

| Carotenoid Pigment | B | **S<sub>R</sub>** | **S<sub>Y</sub>** | **S<sub>G</sub>** | **S<sub>B</sub>** | **S<sub>V</sub>** |
| --- | --- | --- | --- | --- | --- | --- |
| Apocarotenoid | ns | + ** | ns | + *** | ns | ns |
| Canary Xanthophyll Ester 1 | + * | + ** | ns | ns | - * | - * |
| Canary Xanthophyll Ester 2 | + * | + ** | ns | ns | - * | - * |
| Canary Xanthophyll Ester 3 | ns | + ** | ns | ns | - * | - * |
| Echinenone | ns | ns | ns | ns | ns | ns |
| Canthaxanthin Ester 1 | ns | + * | ns | ns | ns | ns |
| Canthaxanthin Ester 2 | ns | + * | ns | ns | ns | ns |
| β-carotene | + *** | + *** | ns | ns | - ** | - ** |
| Ketocarotenoid Ester 2 | + * | + * | + * | ns | - * | - * |
| Cis Lutein Ester | + *** | + *** | ns | ns | - ** | - ** |
| Lutein Ester 1 | + ** | + ** | ns | ns | - * | - * |

**Sup Table 7.** Kit Nonsynonymous SNPs. HS: Hydrophobicity Scale: Nonpolar → Polar distributions of amino acids chains, pH7 (kcal/mol)

| Database | Organism | Protein Sequence | HS | Gen Seq | Conditions | Variant identifier |
| --- | --- | --- | --- | --- | --- | --- |
| this study | *O.pumilio* | Val533Ile | 4.04 ><br>4.92 | C>T |  |  |
| ClinVar | Human | Ile542Met | 4.92 ><br>2.35 | T>G | Gastrointestinal stromal tumor;<br>Hereditary cancer-predisposing syndrome | NM_000222.3(KI T):c.1626T>G (p.Ile542Met) |
| ClinVar | Human | Ile542Phe | 4.92 ><br>2.98 | A>T | Gastrointestinal stromal tumor;<br>Hereditary cancer-predisposing syndrome | NM_000222.3(KI T):c.1624A>T (p.Ile542Phe) |
| this study | *O.pumilio* | Thr957 | -2.57 |  |  |  |
| ClinVar | Human | Val972Met | 4.04 ><br>2.35 | G>A | Gastrointestinal stromal tumor;<br>Hereditary cancer-predisposing syndrome | NM_000222.3(KI T):c.2914G>A (p.Val972Met) |
| ClinVar | Human | Val972Ala | 4.04 ><br>1.81 | T>C | Gastrointestinal stromal tumor | NM_000222.3(KI T):c.2915T>C (p.Val972Ala) |
| this study | *O.pumilio* | Ser958 | -3.4 |  |  |  |
| ClinVar | Human | His973= | -4.66 | C>T | Gastrointestinal stromal tumor;<br>Hereditary cancer-predisposing syndrome | NM_000222.3(KI T):c.2919C>T (p.His973=) |
| this study | *O.pumilio* | Gly959Arg | 0.94 ><br>-14.92 | C>T |  |  |
| ClinVar | Human | Asp974Glu | -8.72 ><br>-6.81 | C>G | Gastrointestinal stromal tumor | NM_000222.3(KI T):c.2922C>G (p.Asp974Glu) |
| ClinVar | Human | Asp974Gly | -8.72 ><br>0.94 | A>G | Gastrointestinal stromal tumor | NM_000222.3(KI T):c.2921A>G (p.Asp974Gly) |
| ClinVar | Human | Asp974Asn | -8.72 > | G>A | Gastrointestinal stromal tumor;<br>Hereditary cancer-predisposing syndrome | NM_000222.3(KI T):c.2920G>A (p.Asp974Asn) |
| this study | *O.pumilio* | Glu960 | -6.81 |  |  |  |
| ClinVar | Human | Asp975His | -8.72 ><br>-4.66 | G>C | Mastocytosis<br>Gastrointestinal stromal tumor<br>**Piebaldism : refers to the absence of mature melanin-forming cells** **(melanocytes) in certain areas of the skin and hair** | NM_000222.3(KI T):c.2923G>C (p.Asp975His) |
| ClinVar | Human | Asp975Asn | -8.72 > -6.64 | G>A | Gastrointestinal stromal tumor; Hereditary cancer-predisposing syndrome | NM_000222.3(KIT):c.2923G>A (p.Asp975Asn) |
| ClinVar | Human | Asp975Gly | -8.72 > 0.94 | A>G | Gastrointestinal stromal tumor; Hereditary cancer-predisposing syndrome | NM_000222.3(KIT):c.2924A>G (p.Asp975Gly) |
| ClinVar | Human | Asp975Glu | -8.72 > -6.81 | T>A | Hereditary cancer-predisposing syndrome | NM_000222.3(KIT):c.2925T>A (p.Asp975Glu) |
| this study | *O.pumilio* | Arg961 | -14.92 |  |  |  |
| ClinVar | Human | Val976Ile | 4.04 > 4.92 | G>A | Gastrointestinal stromal tumor | NM_000222.3(KIT):c.2926G>A (p.Val976Ile) |

## References

1. Daly, J. W. & Myers, C. W. Toxicity of Panamanian poison frogs (Dendrobates): some biological and chemical aspects. Science 156, 970–973 (1967).

2. Coleman, J. L. Why the Striking Diversity of Conspicuous Color Patterns in a Poison Frog from Central America? 58, 70–74 (2023).

3. Summers, K., Bermingham, E., Weigt, L., McCafferty, S. & Dahlstrom, L. Phenotypic and genetic divergence in three species of dart-poison frogs with contrasting parental behavior. J. Hered. 88, 8–13 (1997).

4. Anderson, R. P. & Handley, C. O., Jr. Dwarfism in insular sloths: biogeography, selection, and evolutionary rate. Evolution 56, 1045–1058 (2002).

5. Summers, K., Cronin, T. W. & Kennedy, T. Variation in spectral reflectance among populations of Dendrobates pumilio , the strawberry poison frog, in the Bocas del Toro Archipelago, Panama. J. Biogeogr. 30, 35–53 (2003).

6. Yang, Y., Servedio, M. R. & Richards-Zawacki, C. L. Imprinting sets the stage for speciation. Nature 574, 99–102 (2019).

7. Dreher, C. E., Cummings, M. E. & Pröhl, H. An Analysis of Predator Selection to Affect Aposematic Coloration in a Poison Frog Species. PLoS One 10, e0130571 (2015).

8. Prates, I., Paz, A., Brown, J. L. & Carnaval, A. C. Effects of prey turnover on poison frog toxins: using landscape ecology to assess how biotic interactions affect species phenotypes. bioRxiv 695171 (2019) doi:10.1101/695171.

9. Dugas, M. B., Halbrook, S. R., Killius, A. M., del Sol, J. F. & Richards-Zawacki, C. L. Colour and escape behaviour in polymorphic populations of an aposematic poison frog. Ethology 121, 813–822 (2015).

10. Rudh, A., Rogell, B. & Höglund, J. Non-gradual variation in colour morphs of the strawberry poison frog Dendrobates pumilio: genetic and geographical isolation suggest a role for selection in maintaining polymorphism. Mol. Ecol. 16, 4284–4294 (2007).

11. Pröhl, H. & Hödl, W. Parental investment, potential reproductive rates, and mating system in the strawberry dart-poison frog, Dendrobates pumilio. Behav. Ecol. Sociobiol. 46, 215–220 (1999).

12. Maan, M. E. & Cummings, M. E. Poison frog colors are honest signals of toxicity, particularly for bird predators. Am. Nat. 179, E1–14 (2012).

13. Yang, Y., Dugas, M. B., Sudekum, H. J., Murphy, S. N. & Richards-Zawacki, C. L. Male-male aggression is unlikely to stabilize a poison frog polymorphism. J. Evol. Biol. 31, 457–468 (2018).

14. Richards-Zawacki, C. L., Wang, I. J. & Summers, K. Mate choice and the genetic basis for colour variation in a polymorphic dart frog: inferences from a wild pedigree. Mol. Ecol. 21, 3879–3892 (2012).

15. Richards-Zawacki, C. L. & Cummings, M. E. Intraspecific reproductive character displacement in a polymorphic poison dart frog, Dendrobates pumilio. Evolution 65, 259–267 (2011).

16. Dugas, M. B. & Richards-Zawacki, C. L. A captive breeding experiment reveals no evidence of reproductive isolation among lineages of a polytypic poison frog. Biol. J. Linn. Soc. Lond. 116, 52–62 (2015).

17. Wang, I. J. & Summers, K. Genetic structure is correlated with phenotypic divergence rather than geographic isolation in the highly polymorphic strawberry poison-dart frog. Mol. Ecol. 19, 447–458 (2010).

18. Ruxton, G. D., Allen, W. L., Sherratt, T. N. & Speed, M. P. Avoiding Attack: The Evolutionary Ecology of Crypsis, Aposematism, and Mimicry. (Oxford University Press, 2019).

19. Noonan, B. P. & Comeault, A. A. The role of predator selection on polymorphic aposematic poison frogs. Biol. Lett. 5, 51–54 (2009).

20. Yang, Y., Blomenkamp, S., Dugas, M. B., Richards-Zawacki, C. L. & Pröhl, H. Mate Choice versus Mate Preference: Inferences about Color-Assortative Mating Differ between Field and Lab Assays of Poison Frog Behavior. Am. Nat. 193, 598–607 (2019).

21. Gehara, M., Summers, K. & Brown, J. L. Population expansion, isolation and selection: novel insights on the evolution of color diversity in the strawberry poison frog. Evol. Ecol. 27, 797–824 (2013).

22. Tazzyman, S. J. & Iwasa, Y. SEXUAL SELECTION CAN INCREASE THE EFFECT OF RANDOM GENETIC DRIFT—A QUANTITATIVE GENETIC MODEL OF POLYMORPHISM IN OOPHAGA PUMILIO, THE STRAWBERRY POISON-DART FROG. Evolution 64, 1719–1728 (2010).

23. Lehman, M. R., González-Santoro, M. & Richards-Zawacki, C. L. Little evidence for color- or size-based mating preferences by male strawberry poison frogs (Oophaga pumilio). Behav. Ecol. Sociobiol. 78, 19 (2024).

24. Summers, K., Symula, R., Clough, M. & Cronin, T. Visual mate choice in poison frogs. Proc. Biol. Sci. 266, 2141–2145 (1999).

25. Reynolds, R. G. & Fitzpatrick, B. M. Assortative mating in poison-dart frogs based on an ecologically important trait. Evolution 61, 2253–2259 (2007).

26. Jin, Y. et al. Population Genomics of Variegated Toad-Headed Lizard Phrynocephalus versicolor and Its Adaptation to the Colorful Sand of the Gobi Desert. Genome Biol. Evol. 14, (2022).

27. Bagnara, J. T., Frost, S. K. & Matsumoto, J. On the development of pigment patterns in amphibians. Am. Zool. 18, 301–312 (1978).

28. Bagnara, J. T., Fernandez, P. J. & Fujii, R. On the blue coloration of vertebrates. Pigment Cell Res. 20, 14–26 (2007).

29. Toomey, M. B. et al. A mechanism for red coloration in vertebrates. Curr. Biol. (2022) doi:10.1016/j.cub.2022.08.013.

30. Bagnara, J. T., Taylor, J. D. & Hadley, M. E. The dermal chromatophore unit. J. Cell Biol. 38, 67–79 (1968).

31. Wang, I. J. & Shaffer, H. B. Rapid color evolution in an aposematic species: a phylogenetic analysis of color variation in the strikingly polymorphic strawberry poison-dart frog. Evolution 62, 2742–2759 (2008).

32. Kinoshita, S. & Yoshioka, S. Structural colors in nature: the role of regularity and irregularity in the structure. Chemphyschem 6, 1442–1459 (2005).

33. Ogilvy, V., Preziosi, R. F. & Fidgett, A. L. A brighter future for frogs? The influence of carotenoids on the health, development and reproductive success of the red-eye tree frog. Anim. Conserv. 15, 480–488 (2012).

34. Stückler, S., Cloer, S., Hödl, W. & Preininger, D. Carotenoid intake during early life mediates ontogenetic colour shifts and dynamic colour change during adulthood. Anim. Behav. 187, 121–135 (2022).

35. Taboada, C. et al. Multiple origins of green coloration in frogs mediated by a novel biliverdin-binding serpin. Proc. Natl. Acad. Sci. U. S. A. 117, 18574–18581 (2020).

36. Silverstone, P. A. A Revision of the Poison-Arrow Frogs of the Genus Phyllobates Bibron in Sagra (family Dendrobatidae). (Natural History Museum of Los Angeles County, 1976).

37. Hauswaldt, J. S., Ludewig, A.-K., Vences, M. & Pröhl, H. Widespread co-occurrence of divergent mitochondrial haplotype lineages in a Central American species of poison frog (Oophaga pumilio). J. Biogeogr. 38, 711–726 (2011).

38. Rodríguez, A., Mundy, N. I., Ibáñez, R. & Pröhl, H. Being red, blue and green: the genetic basis of coloration differences in the strawberry poison frog (Oophaga pumilio). BMC Genomics 21, 301 (2020).

39. Stuckert, A. M. M. et al. Transcriptomic analyses during development reveal mechanisms of integument structuring and color production. Evol. Ecol. (2023) doi:10.1007/s10682-023-10256-2.

40. Freeborn, L. R. The Genetic, Cellular, and Evolutionary Basis of Skin Coloration in the Highly Polymorphic Poison Frog, Oophaga pumilio. (University of Pittsburgh, Ann Arbor, United States, 2020).

41. Rogers, R. L. et al. Genomic Takeover by Transposable Elements in the Strawberry Poison Frog. Mol. Biol. Evol. 35, 2913–2927 (2018).

42. Cheng, J. Y., Mailund, T. & Nielsen, R. Fast admixture analysis and population tree estimation for SNP and NGS data. Bioinformatics 33, 2148–2155 (2017).

43. Nielsen, S. V. et al. Bayesian inference of admixture graphs on Native American and Arctic populations. PLoS Genet. 19, e1010410 (2023).

44. Gutenkunst, R., Hernandez, R., Williamson, S. & Bustamante, C. Diffusion Approximations for Demographic Inference: DaDi. Nature Precedings 1–1 (2010).

45. Crawford, A. J. Relative rates of nucleotide substitution in frogs. J. Mol. Evol. 57, 636–641 (2003).

46. Santos, J. C. Fast molecular evolution associated with high active metabolic rates in poison frogs. Mol. Biol. Evol. 29, 2001–2018 (2012).

47. Dessinioti, C., Stratigos, A. J., Rigopoulos, D. & Katsambas, A. D. A review of genetic disorders of hypopigmentation: lessons learned from the biology of melanocytes. Exp. Dermatol. 18, 741–749 (2009).

48. Rohrlich, S. T. & Porter, K. R. Fine structural observations relating to the production of color by the iridophores of a lizard. Anolis carolinensis. J. Cell Biol. 53, 38–52 (1972).

49. Smyth, I. M. et al. Genomic anatomy of the Tyrp1 (brown) deletion complex. Proc. Natl. Acad. Sci. U. S. A. 103, 3704–3709 (2006).

50. Fang, W., Huang, J., Li, S. & Lu, J. Identification of pigment genes (melanin, carotenoid and pteridine) associated with skin color variant in red tilapia using transcriptome analysis. Aquaculture 547, 737429 (2022).

51. Hooper, D. M., Griffith, S. C. & Price, T. D. Sex chromosome inversions enforce reproductive isolation across an avian hybrid zone. Mol. Ecol. 28, 1246–1262 (2019).

52. Ahi, E. P. et al. Comparative transcriptomics reveals candidate carotenoid color genes in an East African cichlid fish. BMC Genomics 21, 54 (2020).

53. McKinnon, J. S., Newsome, W. B. & Balakrishnan, C. N. Gene expression in male and female stickleback from populations with convergent and divergent throat coloration. Ecol. Evol. 12, e8860 (2022).

54. Stuckert, A. M. M. et al. The genomics of mimicry: Gene expression throughout development provides insights into convergent and divergent phenotypes in a Müllerian mimicry system. Mol. Ecol. 30, 4039–4061 (2021).

55. Moest, M. et al. Selective sweeps on novel and introgressed variation shape mimicry loci in a butterfly adaptive radiation. PLoS Biol. 18, e3000597 (2020).

56. von Lintig, J. & Vogt, K. Filling the Gap in Vitamin A Research: MOLECULAR IDENTIFICATION OF AN ENZYME CLEAVING β-CAROTENE TO RETINAL *. J. Biol. Chem. 275, 11915–11920 (2000).

57. Kloer, D. P. & Schulz, G. E. Structural and biological aspects of carotenoid cleavage. Cell. Mol. Life Sci. 63, 2291–2303 (2006).

58. Cortés, R. et al. Evolution of the melanocortin system. Gen. Comp. Endocrinol. 209, 3–10 (2014).

59. Kobayashi, Y., Tsuchiya, K., Yamanome, T., Schiöth, H. B. & Takahashi, A. Differential expressions of melanocortin receptor subtypes in melanophores and xanthophores of barfin flounder. Gen. Comp. Endocrinol. 168, 133–142 (2010).

60. Mizusawa, K., Yamamura, Y., Kasagi, S., Cerdá-Reverter, J. M. & Takahashi, A. Expression of genes for melanotropic peptides and their receptors for morphological color change in goldfish Carassius auratus. Gen. Comp. Endocrinol. 264, 138–150 (2018).

61. Han, J., Hong, W. S., Wang, Q., Zhang, T. T. & Chen, S. X. The regulation of melanocyte-stimulating hormone on the pigment granule dispersion in the xanthophores and melanophores of the large yellow croaker (Larimichthys crocea). Aquaculture 507, 7–20 (2019).

62. Kobayashi, Y., Mizusawa, K., Chiba, H., Tagawa, M. & Takahashi, A. Further evidence on acetylation-induced inhibition of the pigment-dispersing activity of α-melanocyte-stimulating hormone. Gen. Comp. Endocrinol. 176, 9–17 (2012).

63. Braasch, I., Brunet, F., Volff, J.-N. & Schartl, M. Pigmentation pathway evolution after whole-genome duplication in fish. Genome Biol. Evol. 1, 479–493 (2009).

64. Samaniego Castruita, J. A., Westbury, M. V. & Lorenzen, E. D. Analyses of key genes involved in Arctic adaptation in polar bears suggest selection on both standing variation and de novo mutations played an important role. BMC Genomics 21, 543 (2020).

65. I P Tomlinson, R. S. H. Peutz-Jeghers syndrome. J. Med. Genet. 34, 1007 (1997).

66. Chae, H.-D. & Jeon, C.-H. Peutz-Jeghers syndrome with germline mutation of STK11. Ann Surg Treat Res 86, 325–330 (2014).

67. Korsse, S. E., van Leerdam, M. E. & Dekker, E. Gastrointestinal diseases and their oro-dental manifestations: Part 4: Peutz-Jeghers syndrome. Br. Dent. J. 222, 214–217 (2017).

68. Shevde, L. A. et al. Suppression of human melanoma metastasis by the metastasis suppressor gene, BRMS1. Exp. Cell Res. 273, 229–239 (2002).

69. Maeda, A. et al. Redundant and unique roles of retinol dehydrogenases in the mouse retina. Proc. Natl. Acad. Sci. U. S. A. 104, 19565–19570 (2007).

70. Stuckert, A. M. M. et al. Variation in pigmentation gene expression is associated with distinct aposematic color morphs in the poison frog Dendrobates auratus. BMC Evol. Biol. 19, 85 (2019).

71. Chovnick, A., Gelbart, W. & McCarron, M. Organization of the Rosy locus in Drosophila melanogaster. Cell 11, 1–10 (1977).

72. Rubio, A. O., Stuckert, A. M. M., LaPolice, T. M., Cole, T. J. & Summers, K. Under pressure: evidence for selection on color-related genes in poison frogs of the genus Ranitomeya. Evol. Ecol. (2024) doi:10.1007/s10682-024-10297-1.

73. Yuzawa, S. et al. Structural basis for activation of the receptor tyrosine kinase KIT by stem cell factor. Cell 130, 323–334 (2007).

74. Hegna, R. H., Saporito, R. A. & Donnelly, M. A. Not all colors are equal: predation and color polytypism in the aposematic poison frog Oophaga pumilio. Evol. Ecol. 27, 831–845 (2013).

75. Galeano, S. P. & Harms, K. E. Coloration in the polymorphic frog Oophaga pumilio associates with level of aggressiveness in intraspecific and interspecific behavioral interactions. Behav. Ecol. Sociobiol. 70, 83–97 (2016).

## References

1. Rodríguez, A., Mundy, N. I., Ibáñez, R. & Pröhl, H. Being red, blue and green: the genetic basis of coloration differences in the strawberry poison frog (Oophaga pumilio). BMC Genomics 21, 301 (2020).

2. Fang, W., Huang, J., Li, S. & Lu, J. Identification of pigment genes (melanin, carotenoid and pteridine) associated with skin color variant in red tilapia using transcriptome analysis. Aquaculture 547, 737429 (2022).

3. Toomey, M. B. et al. High-density lipoprotein receptor SCARB1 is required for carotenoid coloration in birds. Proc. Natl. Acad. Sci. U. S. A. 114, 5219–5224 (2017).

4. Stuckert, A. M. M. et al. The genomics of mimicry: Gene expression throughout development provides insights into convergent and divergent phenotypes in a Müllerian mimicry system. Mol. Ecol. 30, 4039–4061 (2021).

5. Yang, Z. et al. Genetic adaptation of skin pigmentation in highland Tibetans. Proc. Natl. Acad. Sci. U. S. A. 119, e2200421119 (2022).

6. Liu, S. et al. Population genomics reveal recent speciation and rapid evolutionary adaptation in polar bears. Cell 157, 785–794 (2014).

7. Kunieda, T., Nakagiri, M., Takami, M., Ide, H. & Ogawa, H. Cloning of bovine LYST gene and identification of a missense mutation associated with Chediak-Higashi syndrome of cattle. Mamm. Genome 10, 1146–1149 (1999).

8. Anistoroaei, R., Krogh, A. K. & Christensen, K. A frameshift mutation in theLYSTgene is responsible for the Aleutian color and the associated Chédiak-Higashi syndrome in American mink. Anim. Genet. 44, 178–183 (2013).

9. Ullate-Agote, A. et al. Genome mapping of a LYST mutation in corn snakes indicates that vertebrate chromatophore vesicles are lysosome-related organelles. Proc. Natl. Acad. Sci. U. S. A. 117, 26307–26317 (2020).

10. Tharmarajah, G. T. The role of Adamts9 in melanoblast migration and modification of the skin proteome. Preprint at 10.14288/1.0166255 (2015).

11. McLean, C. A., Lutz, A., Rankin, K. J., Stuart-Fox, D. & Moussalli, A. Revealing the Biochemical and Genetic Basis of Color Variation in a Polymorphic Lizard. Mol. Biol. Evol. 34, 1924–1935 (2017).

12. Kelsh, R. N., Harris, M. L., Colanesi, S. & Erickson, C. A. Stripes and belly-spots -- a review of pigment cell morphogenesis in vertebrates. Semin. Cell Dev. Biol. 20, 90–104 (2009).

13. Braasch, I., Brunet, F., Volff, J.-N. & Schartl, M. Pigmentation pathway evolution after whole-genome duplication in fish. Genome Biol. Evol. 1, 479–493 (2009).

14. Qvarnström, A. et al. Coarse dark patterning functionally constrains adaptive shifts from aposematism to crypsis in strawberry poison frogs. Evolution 68, 2793–2803 (2014).

15. Chen, H. et al. Analysis of recently duplicated TYRP1 genes and their effect on the formation of black patches in Oujiang-color common carp (Cyprinus carpio var. color). Anim. Genet. 52, 451–460 (2021).

16. Jolivet, P. & Joseph. SPRI bead mix v1. protocols.io ZappyLab, Inc. 10.17504/protocols.io.bnz4mf8w (2020).

17. Rogers, R. L. et al. Genomic Takeover by Transposable Elements in the Strawberry Poison Frog. Mol. Biol. Evol. 35, 2913–2927 (2018).

18. Li, H. Aligning sequence reads, clone sequences and assembly contigs with BWA-MEM. arXiv [q-bio.GN] (2013).

19. Danecek, P. et al. Twelve years of SAMtools and BCFtools. Gigascience 10, (2021).

20. Korneliussen, T. S., Albrechtsen, A. & Nielsen, R. ANGSD: Analysis of Next Generation Sequencing Data. BMC Bioinformatics 15, 356 (2014).

21. Meisner, J. & Albrechtsen, A. Inferring Population Structure and Admixture Proportions in Low-Depth NGS Data. Genetics 210, 719–731 (2018).

22. Cheng, J. Y., Mailund, T. & Nielsen, R. Fast admixture analysis and population tree estimation for SNP and NGS data. Bioinformatics 33, 2148–2155 (2017).

23. Gutenkunst, R., Hernandez, R., Williamson, S. & Bustamante, C. Diffusion Approximations for Demographic Inference: DaDi. Nature Precedings 1–1 (2010).

24. Gehara, M., Summers, K. & Brown, J. L. Population expansion, isolation and selection: novel insights on the evolution of color diversity in the strawberry poison frog. Evol. Ecol. 27, 797–824 (2013).

25. van der Walt, S. et al. scikit-image: image processing in Python. PeerJ 2, e453 (2014).

26. Simonyan, K. & Zisserman, A. Very Deep Convolutional Networks for Large-Scale Image Recognition. arXiv [cs.CV] (2014).

27. Hudson, R. R., Kreitman, M. & Aguadé, M. A test of neutral molecular evolution based on nucleotide data. Genetics 116, 153–159 (1987).

28. Kim, S. Y. et al. Estimation of allele frequency and association mapping using next-generation sequencing data. BMC Bioinformatics 12, 231 (2011).

29. Korneliussen, T. S., Moltke, I., Albrechtsen, A. & Nielsen, R. Calculation of Tajima’s D and other neutrality test statistics from low depth next-generation sequencing data. BMC Bioinformatics vol. 14 Preprint at 10.1186/1471-2105-14-289 (2013).

30. Tajima, F. Statistical method for testing the neutral mutation hypothesis by DNA polymorphism. Genetics 123, 585–595 (1989).

